# Intermittent brain network reconfigurations and the resistance to social media influence

**DOI:** 10.1101/2021.12.07.471625

**Authors:** Italo’Ivo Lima Dias Pinto, Nuttida Rungratsameetaweemana, Kristen Flaherty, Aditi Periyannan, Amir Meghdadi, Christian Richard, Chris Berka, Kanika Bansal, Javier Omar Garcia

## Abstract

Since their development, social media has grown as a source of information and has a significant impact on opinion formation. Individuals interact with others and content via social media platforms in a variety of ways but it remains unclear how decision making and associated neural processes are impacted by the online sharing of informational content, from factual to fabricated. Here, we use EEG to estimate dynamic reconfigurations of brain networks and probe the neural changes underlying opinion change (or formation) within individuals interacting with a simulated social media platform. Our findings indicate that the individuals who changed their opinions are characterized by less frequent network reconfigurations while those who did not change their opinions tend to have more flexible brain networks with frequent reconfigurations. The nature of these frequent network configurations suggests a fundamentally different thought process between intervals in which individuals are easily influenced by social media and those in which they are not. We also show that these reconfigurations are distinct to the brain dynamics during an in-person discussion with strangers on the same content. Together, these findings suggest that brain network reconfigurations may not only be diagnostic to the informational context but also the underlying opinion formation.

**Author Summary:** Distinctive neural underpinnings of opinion formation and change during in-person and online social interactions are not well understood. Here, we analyze EEG recordings of the participants interacting with a simulated social media platform and during an in-person discussion using a network-based analysis approach. We show that the structure of network reconfigurations during these interactions is diagnostic of the opinion change and the context in which information was received.

## Introduction

Decision making is the internal process by which information is reduced to a categorical and actionable proposition (for review, see Gold & Shadlen, 2007). In the brain, the decision making process has been described as a non-linear, context-dependent process that requires a variety of brain areas to receive and interpret information (e.g., sensory), establish value of this information, and then, based on prior experience and motivation, use a decision variable to produce the proposition and subsequently *act* (Fellows, 2004). One context that is currently and almost ubiquitously used as a source of information is social media, the suite of interactive online technologies that have become a mainstay of not only our everyday interactions but also current events and global happenings (Westerman et. al., 2014). Because of the ubiquitous nature of social media, the unbridled spread of information through it (Yoo, et. al. 2016), and the potentially negative consequences of it (Keles et. al., 2020), it is important to understand how it shapes our thoughts, influences our opinions, and impacts our future actions.

The neurological processes underlying the formation or changing of opinions due to social media exposure have been studied from the perspective of the presence and nature of biased content, and the way in which others interact with the information (e.g, likes, comments, retweets, etc.). Prior neuroscience work has specifically studied the effect of social influence on opinion formation and opinion change within the social media environment, where a network of brain regions including the striatum, orbitofrontal cortex, and temporoparietal junction appear to have a critical role in this decision making process (Cascio et al., 2015; Casado-Aranda et al., 2020; Sherman et al., 2016; Baek et al.,2021; Nakao et al., 2016; Falk et al., 2012; Falk & Scholz, 2018; Kappes et al., 2020; Izuma & Adolphs, 2013; Li et al., 2019; Klucharev et al., 2011). Specifically, the neural mechanism of opinion change due to *social media use* has been shown to integrate brain areas of the valuation, social pain/exclusion, and mentalizing systems which include the ventro-medial prefrontal cortex (VMPFC), striatum, medial prefrontal cortex (mPFC), dorsomedial prefrontal cortex (DMPFC), temporo-parietal junction (TPJ), posterior cingulate (PCC), medial tegmental gyrus (MTG), and anterior cingulate (ACC) (Cascio et al., 2015; Baek et al.,2021; Kappes et al., 2020; Falk et al., 2012). Other work has suggested that the popularity of content (Sherman et al., 2016) and the valence of the content plays a significant role in swaying opinion on these platforms (Baek et al., 2021). Due to the opportunity social media affords in rapidly disseminating information throughout the globe, it also creates an interesting glimpse into the complex human decision making process that impacts our everyday lives (Schmälzle et. al., 2017). Indeed, with the intensity and speed in which *information* spreads in this media convolved with the global scale, the contextual impact on decisions derived from platforms like these have had demonstrably profound impacts on society as a whole (Spinney, 2017).

Despite the understanding of the importance of these platforms in forming our decisions, it is still unclear how brain networks composed of regions, perhaps those associated with social media informational processing and influence, interact to produce opinion change. Importantly, it is also unclear how this process may be unique to brain processes underlying in-person interaction and free discussion. Network neuroscience provides a variety of tools to understand the complex network properties of the brain and has proven successful in describing a variety of behaviors (e.g., Bassett and Sporns, 2017 or Betzel and Bassett, 2017). For example, the rate at which networks within the brain rapidly reconfigure to support cognition has been found to be highly predictive of a variety of cognitive processes. Dynamic community detection, a technique used to distill complex connectivity patterns into time-varying labels of *communities* (i.e., clusters of nodes) has been successful in capturing variability in a variety of behaviors in fMRI studies primarily, but has also been used for understanding band-specific EEG connectivity patterns (e.g., Garcia et al., 2020). Here, we have investigated the rapid fluctuations in network connectivity while individuals are exposed to an interactive social media platform containing factual and fake content attempting to simulate the real-world experience of social media and relate this to changes in opinions after exposure to this content, as we hypothesize that the flexible dynamics within the brain may be associated with complex decision making behind opinion change. Importantly, we provide a comparison to in-person discussion which allows us to disentangle the unique neural properties of this process. Our results provide preliminary evidence of unique neural features marking the cognitive processes supporting decision-making prompted by digital stimuli on a social media platform.

## Results

We have investigated the neural correlates of complex decision making during online social media and in-person social interactions and assessed opinion change with questionnaires that asked participants for their opinions on several topical issues. Opinions on these topical issues were gathered before and after the interaction with a simulated social media platform and after in-person discussion of the content (Figure 1, see also Richard et. al., 2021). EEG was concurrently collected (see Supplemental Material Figure S1 for electrode montage) during the social media and in-person interactions and was analyzed to understand the rapid reconfigurations in EEG-derived brain connectivity matrices during the complex process of information gathering and opinion change and/or formation. Here, we used dynamic community detection, an algorithm that has previously been shown to successfully capture brain network reconfigurations associated with the variability in human behavior across a variety of tasks. We extended these findings by inspecting the temporal dynamics of node-pair community affiliations and comparing this metric between those that changed their opinions and those that did not across these social interaction conditions.

**Figure 1:**
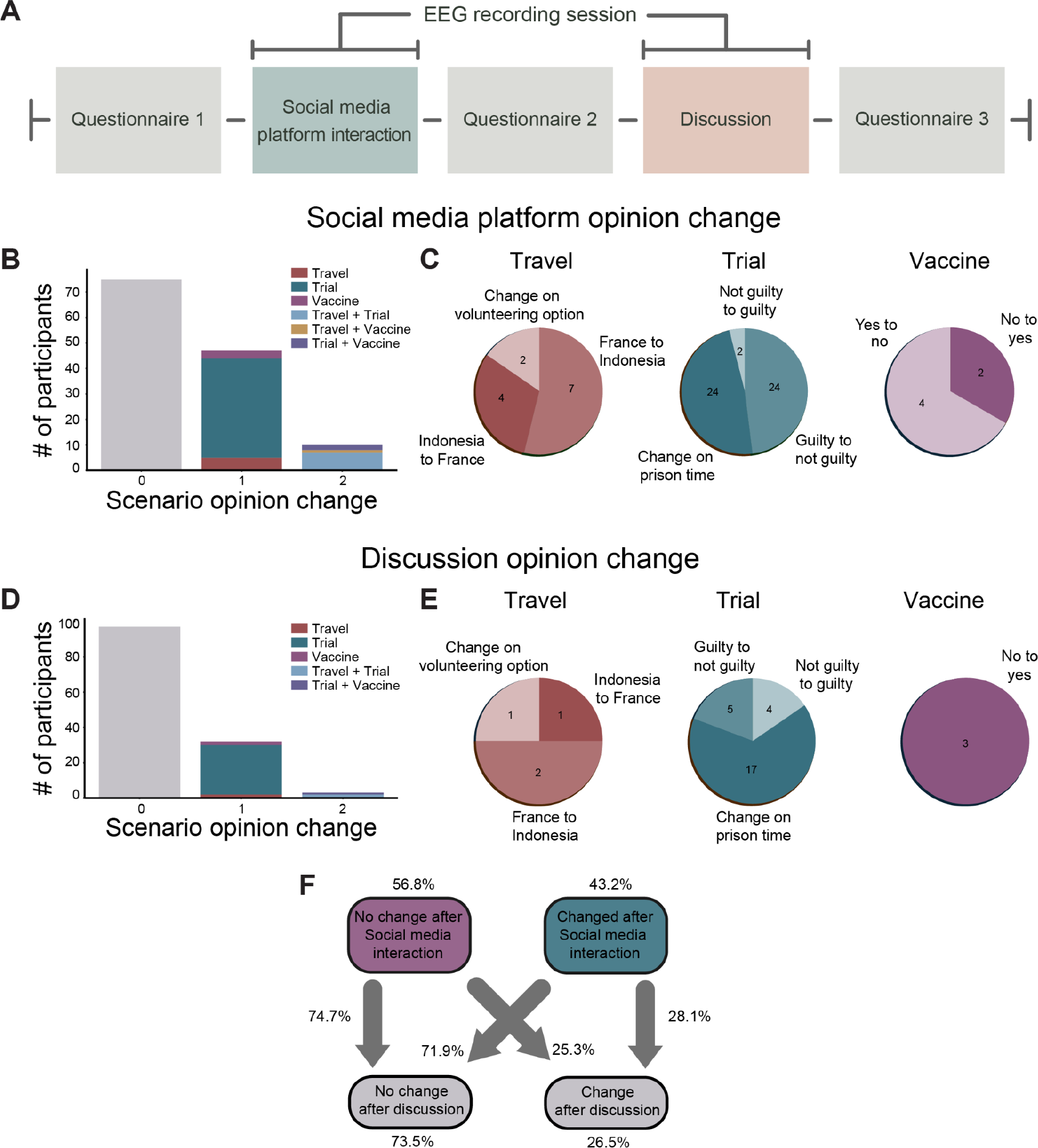
Experimental setup and opinion change quantification. (A) Timeline of the experimental design. The opinion changes of each subject were assessed through the application of a questionnaire before and after the subject interaction with the social media platform, in addition to in-person discussion. (B,D) Histogram of opinion change by scenario in social media interaction and in-person discussion, respectively. 0, 1, and 2 indicate the number of scenarios on which individuals change their opinion. Color legend indicates the scenario. (C,E) Pie charts indicating the direction of opinion change for all of the changes observed after the social media platform and in-person interactions. (F) Flow chart indicating the fraction of participants that changed their opinion from the social media platform (C_s_) interaction to in-person discussion (C_i_).

### Characterizing individuals by opinion change after social media and in-person interactions

Figure 1A shows the experimental timeline, where, after arriving in the laboratory, subjects were presented with questionnaires that asked their opinions on three particular real-world topical issues, namely: (i) travel based on social awareness and volunteerism, (ii) punishment after a murder trial, and (iii) decisions to vaccinate from disease before and after interaction with the social media platform as well as after the in-person discussion segment. EEG was recorded during these two interactive conditions, i.e., *social media interaction* and *in-person discussion*. These interactive conditions differed in several ways. First, during the social media interaction interval, subjects were seated in front of a monitor and were allowed to freely scroll through the simulated social media platform and interact (e.g., “like”, “share” posts) for no more than 2 hours. During the in-person discussion segment, subjects (3-4 at a time) were seated in another room where an experimenter moderated the conversation and asked subjects to discuss the topics for no more than 20 minutes. Based on changes of the questionnaire answers before and after each interaction we grouped the intervals into two segments, those intervals in which subjects changed their opinions (“change”, C) and those that did not (“no change”, NC), and, for shorthand, we use the acronyms C_s_ and NC_s_ for groups that change and did not change their opinion after social media interaction. For in-person discussion, we similarly use the shorthand C_i_ and NC_i_. Figure 1B-D shows the distribution of responses and the rate of opinion change across participants (N = 132) following both conditions. First, with the social media platform, a majority of individuals did not change their opinion from the initial survey (N = 75); however, a total of 57 individuals changed their opinions, with the most individuals changing their opinion in the murder trial scenario (N = 39). A small proportion of the individuals (N = 10) changed their opinions in two scenarios and were most likely to change their opinion on the travel and murder scenarios (Figure 1B). Figure 1C displays a more granular visualization of responses for each scenario, and similar to the design of the experiment which presented equally positive and negative coverage on an issue, there was a large diversity in opinion changes, validating the well-balanced affectual information within the platform. For example, even with the vaccination scenario, there were some individuals who changed their opinion toward not vaccinating after the social media interaction. Importantly, as well, is the fact that the change in prison time in the murder trial contributed to the largest changes, with 24 subjects changing the prison time after the social media interaction and 24 changing from guilty to not guilty. This was significant for both the social media interaction (χ^2^(1, N =132)=7.75, p = .005) and in-person discussion segments (χ^2^(1, N =132)=48.48, p < .0001).

We observed some similarities and differences in opinion change after in-person discussion. As shown in Figure 1D-E, the overwhelming majority of individuals (73.5%, N = 97) did not change their opinion, suggesting that the in-person discussion was less likely to affect one’s opinion than social media interaction; however, the order of the questionnaires was the same across all individuals. This limitation does not allow us to disentangle the effect questionnaire-order may have on our effects. Similar to the social media interaction condition, though, changes in opinion mostly occurred within the murder trial condition, accounting for 80% of the total changes in opinion after the discussion. Very few individuals changed their opinion in more than one scenario, accounting for only 8.6% of the total opinion change after the discussion.

Finally, to characterize the opinion change, overall, we estimated transition probabilities as presented in Figure 1F. These transition probabilities can give us a glimpse into the individual subjects that *did and did not* change their opinions following the social media interaction and in-person interactive conditions. As is shown in Figure 1F, 56.8% of the subjects did not change their responses after social media interaction, 74.7% of these individuals also kept their responses after in-person discussion. From the 43.2% of the subjects that changed their responses after the social media interaction, 28.1% changed their responses after in-person discussion as well. Importantly, of all the participants, 73.5% did not change their opinion after in-person discussion. Due to the distribution of those that did not change their opinion and those that did, we next compared groups of subjects who *did and did not change* their opinions. Critically, if an individual changed their opinion in any of the scenarios, they were included in the “change” (C) group and only those who did not change their opinion in any scenario were included in the “no change” group (NC) for all subsequent analyses.

### Nodal Flexibility distinguishes individuals who changed their opinion following social media interaction

We hypothesized that complex decision making and information processing requires the reconfiguration of underlying brain networks. To test this hypothesis, we applied a dynamic community detection analysis to the EEG data and probed how network reconfigurations are associated with opinion change by directly comparing the intervals in which subjects did and did not change their opinion (Figure 2). This was accomplished in several steps. First, the dynamic community structure requires an estimate of the underlying statistical dependency between nodes. Here, we estimated this statistical dependence, or functional connectivity of the EEG, using the pairwise weighted phase lag index (wPLI) separately for commonly studied EEG oscillations (i.e., delta (1-3 Hz), theta (3-7 Hz), alpha (8-13 Hz), beta (21-30), and gamma (25-40) bands) in non-overlapping 10 sec time windows. This data-driven approach to functional connectivity exploits phase-based relationships within the data, yielding connectivity matrices that are reliable and less susceptible to some expected artifacts without requiring parameterization (Hardmeier et. al., 2014). Once calculated, the wPLI matrices were used to determine the community structure by modularity maximization using a Louvain algorithm (see Methods). This distilled the connectivity time evolving matrices into an average of 315 (SD = 102) time windows of community labels that represent the band-specific community affiliations of EEG sensors across time.

**Figure 2:**
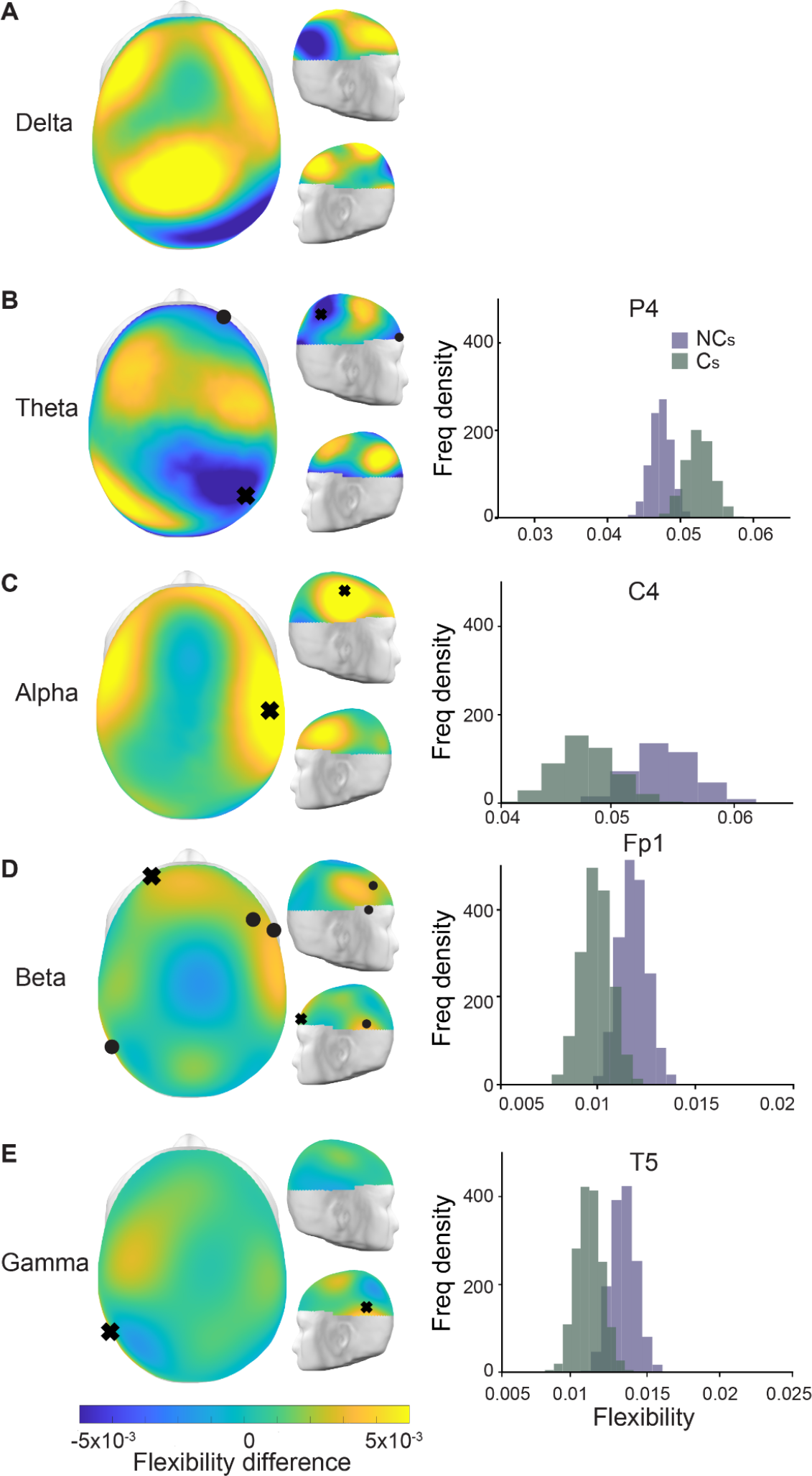
Flexibility differences between the intervals in which individuals did and did not change their opinion. (A-E) Topographic plots of each frequency band showing the difference of the mean flexibility between the two groups such that positive values (in yellow) indicate an increased flexibility for those that did not change their opinions. The black tokens (orbs, x’s) indicate sensors with statistically significant differences in flexibility between the two groups as found via a bootstrap procedure (see Methods). On the right, we show representative bootstrap distributions of the mean flexibility of the sensors marked by an x for the two intervals in purple (NC, no change) and green (C, change).

From these affiliations (i.e., distilled connectivity matrices), we estimated sensor (node) *flexibility,* which is a measure of how much each node changes its affiliation across time between the opinion change groups. Since the data is not balanced between the groups, we employed a bootstrap procedure to estimate the distributions of mean flexibility for each of the EEG sensors for both groups (“change vs no change”, see Methods), and subsequently compared the node flexibility when individuals changed their opinion (or did not). We observed that those intervals in which individuals did not change their opinion showed significant increases in node flexibility in the alpha, beta, and gamma bands and a decrease in flexibility in the theta band. Figure 2 shows the node flexibility differences, yellow (and blue) shades indicate an increased flexibility on the group without change (and with change). An increased node flexibility in those with no change in opinions was observed in the higher frequency bands, with beta band flexibility showing significant differences (bootstrap analysis, *p* < 0.05) for sensors F8, F4, Fp1 and T5, T5 in the gamma band, and C4 in the alpha band. In contrast, the theta band presents a statistically significant decrease in node flexibility in those that did not change their opinion compared to those who did sensors P4 and Fp2, differing from the other frequency bands. These results indicate that the dynamics of the synchronization-desynchronization processes, as measured by the wPLI, plays an important role in the underlying mechanism of opinion change whilst interacting with social media.

### Assessing dynamic changes in community structure

We found that the node flexibility is informative as a neural marker of opinion change; however, it does not provide much information of the dynamic changes in the community structure to further understand the underlying network reconfigurations leading to opinion change. For example, one could ask how the links of the flexible nodes evolve with time and which other nodes couple and decouple with them more often during the task. In this regard, allegiance is a commonly used metric that captures the fraction of time two nodes share the same community affiliation, 0 for a pair of nodes that never share a community and 1 for nodes that are always in the same community. We estimated node allegiances for individuals with and without a change in opinion and found that they do not differentiate the two groups (for further details, see Supplemental Materials and Figure S2). However, it is also unclear from allegiances alone whether more fine-grained temporal dynamics of network reconfigurations might differentiate these groups. To more finely understand the temporal evolution of node-pair affiliation change, we computed a new metric called *intermittence*.

Like allegiance, intermittence is a measure of the interaction between two nodes of the network; however, while allegiance captures the fraction of time two nodes belong to the same community, intermittence tracks how frequently the two nodes change their affiliation from the same to different and vice versa. In other words, intermittence differentiates two nodes’ affiliation changes that occur in rapid bursts from affiliation changes that occur in longer-term after more static community affiliation similarity. Together, we may inspect allegiance as the likelihood for two nodes to be in the same community, and intermittence can inform us of the temporal nature of this relationship.

### Intermittence differentiates changes in opinion

In exploring the intermittence metric, we first directly compare the allegiance and intermittence metrics for both groups (Figure 3A). We observe for lower frequency bands (e.g., delta) that intermittence is more variable, spanning a wider range of values than higher frequency bands while the opposite is true for allegiance. Specifically for the delta band, values of allegiance larger than 0.6 are less frequent than observed for the other frequency bands and values of intermittence above 0.1 are more frequent than for the beta and gamma bands. On the other hand, inspecting higher frequency oscillations, we see intermittence is rarely above 0.05 but allegiance spans the entire range of possible values. This suggests that there is a higher propensity for more static network reconfigurations at higher frequencies than lower frequencies (e.g., compare Figure 3A gamma and delta). Importantly, with intermittence estimation, simply by its calculation, allegiance imposes an inherent restriction on its range of possible values. The maximum value of intermittence for a given pair of nodes is limited by the value of allegiance between those nodes, as the reader should understand that there cannot be more dynamic changes between nodes if they are rarely ever in the same community. Thus, given our observation that higher frequency bands (beta, gamma) had higher allegiance (that could allow for higher intermittence) in addition to the observed lower average intermittence, the finding that higher frequency bands show are even more striking, suggesting higher frequencies display very static network dynamics across groups.

**Figure 3:**
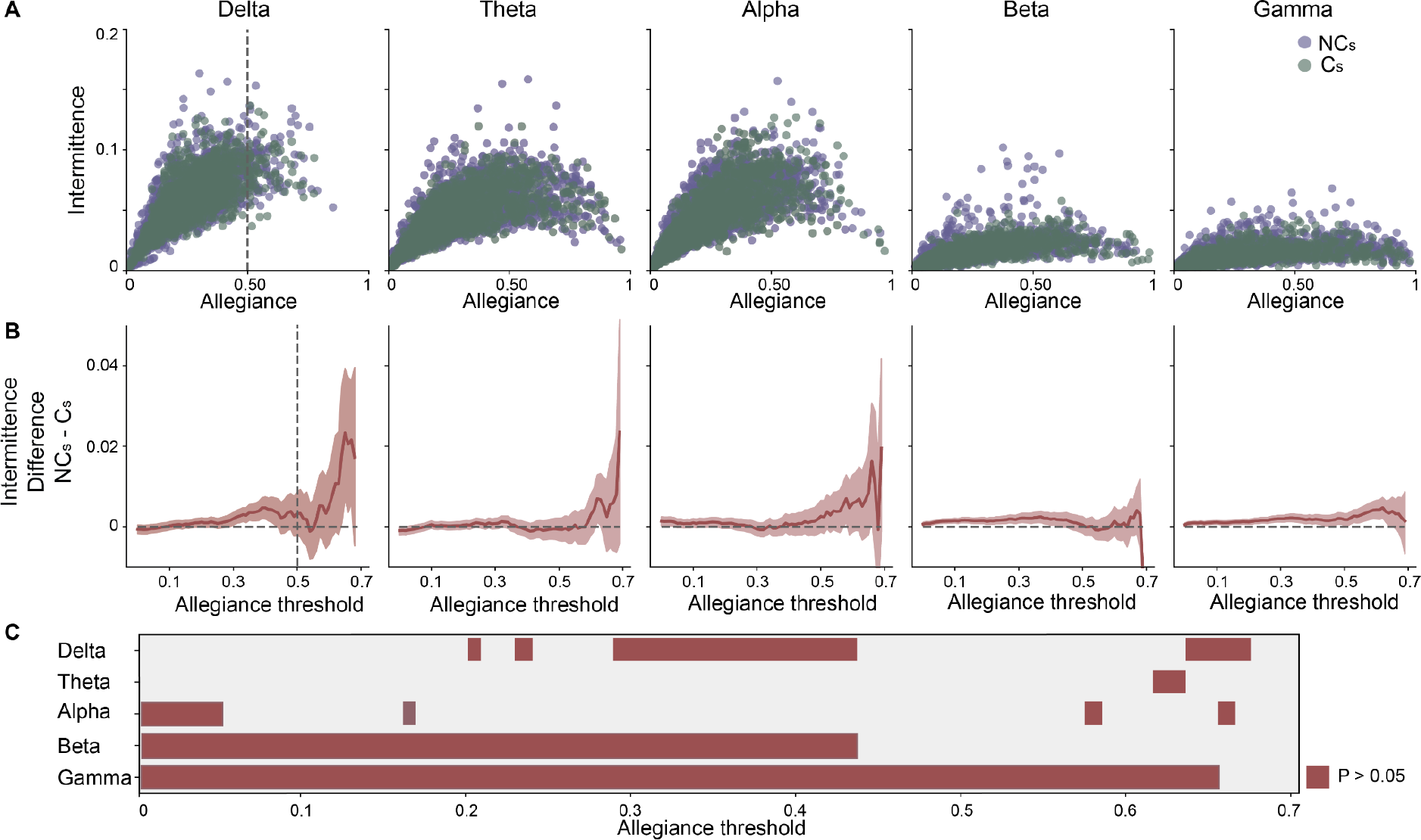
Comparing intermittence and allegiance in opinion change. (A) Scatter plots of the relationship between intermittence and allegiance for each frequency band of interest, where purple (or green) indicates channels of individuals who did not (and did) change their opinion (C vs NC). Dashed vertical lines indicate the middle value of allegiance, to visually anchor the plots. (B) Bootstrapped difference (*no change*-*change; NC-C*) plots of intermittence of the intervals for different threshold levels of allegiance for each frequency band of interest. Shaded region is 95% confidence interval and the vertical dashed line indicates the midpoint of allegiance to visually anchor the midpoint. (C) Allegiance thresholds that survive statistical comparison of opinion change for each frequency band.

In Figure 3B, we show the differences between the means in intermittence of the two groups. Critically, mean bootstrap distributions were calculated using *only those* points with associated allegiance values higher than the allegiance threshold indicated on the x-axis. The shaded area in Figure 3B represents a 95% confidence interval and was obtained by a bootstrap procedure with 10,000 samples (see Methods). To summarize the statistical comparison between those intervals with or without opinion change at each of the frequency bands, Figure 3C displays those allegiance thresholds that display the statistical difference (*p* < 0. 05).

Comparison of the mean intermittence between the two groups shows that the intermittence metric can delineate between those in which there was or was not a change in opinion in each frequency band, but to a highly variable extent. For example, our results show that within the delta band, we observed statistically significant differences for the allegiance threshold range between 0.29 and 0.43 and a few other allegiance values accounting for more than 14% of the possible allegiance range values. For the theta band, there were minimal differences between groups observed, accounting for only 2.8% of the total allegiance range. Within the alpha band, we observed, again, minimal differences between the groups accounting for less than 5% of allegiance thresholds; importantly, they were observed mostly at the lowest allegiance thresholds. The most robust differences between opinion change groups were observed within the beta and gamma bands. For the beta band, we observed significant differences between the groups in approximately 43% of the allegiance threshold range. The lowest p-values we observed between those without a change in opinion (M = 0.018 *a.u.*) and those with a change in opinion (M = 0.017 *a.u.*) was for the allegiance threshold of 0.12 (*p* = 2. 7 × 10^−5^). The largest range of allegiance values with significant differences between the two groups were observed in the gamma band, accounting for nearly 65% of the entire allegiance range. The lowest p-value was observed at an allegiance threshold of 0.29 for no opinion change (M = 0.018 *a.u.*) and opinion change (M = 0.016 *a.u.p* = 3. 6 × 10^−6^) intervals. Thus, it appears that intermittence successfully delineates those individuals with and without a change in opinion whilst interacting with social media, but does so in a frequency band-specific manner where the beta and gamma bands show the most robust differences as indicated by a wide range of allegiance thresholds for which the two groups have a significant difference in intermittence. In other words, intermittence can be used to characterize opinion change in band-specific oscillatory schemes, but it is still unclear whether this is a general opinion change phenomenon or if this may be specific to the context in which information is received (i.e., social media platform). Thus, we next explored how these findings could differ in a different context, specifically during *in-person discussion*.

### Social media interaction and in-person discussion differences

To determine the specificity of our findings to the social interaction context (e.g., social media vs in-person discussion), we sought to determine if network dynamics between these opinion change groups have similar structure during social media and in-person interactions. The in-person discussion was completed after the social media interaction and was conducted by an experimenter who acted as a moderator and prompted individuals to discuss the topics which were probed by the surveys (see Methods). To estimate the difference in neural dynamics between these two social interaction contexts, we calculated the coefficient of variation (CoV) of the time series of the wPLI during social media interaction *and* in-person discussion. This procedure is a computationally inexpensive and complementary approach to capture dynamic reconfiguration of the synchronous patterns estimated from the EEG recordings that we expect to be similarly sensitive to the intermittence metric but on the node-level, like flexibility.

Similar to the EEG measurements during the social media interaction, the statistical dependencies between nodes were estimated with the wPLI metric, and then, in a pairwise fashion, the temporal coefficient of variation (temporal CoV) was calculated for each of the node-pairs. Finally, to aggregate the data for each subject, the mean temporal CoV for each pair was estimated, and for each subject, we calculated the mean across all the nodes. This procedure results in a single mean temporal CoV for each session and finally the group distribution and statistical comparisons were completed with a bootstrap procedure for each social context and opinion change (or no change) and are summarized in Figure 4. We observed clear group differences in the gamma band (p = 0.031) during the social media interaction, in accordance with the results obtained from the intermittence analysis. For those subjects who were more likely to change their opinion, we also observed a difference in context, where we observed more variability (temporal CoV) in the social media interaction than in the in-person discussion (p < 0.05). Moreover, within the delta and alpha bands, we observed this context effect, too, where temporal CoV of connectivity in the social media intervals was significantly higher than in the in-person interaction contexts. Interestingly and divergent from the previous findings, we observed no significant differences in the beta and theta bands.

**Figure 4:**
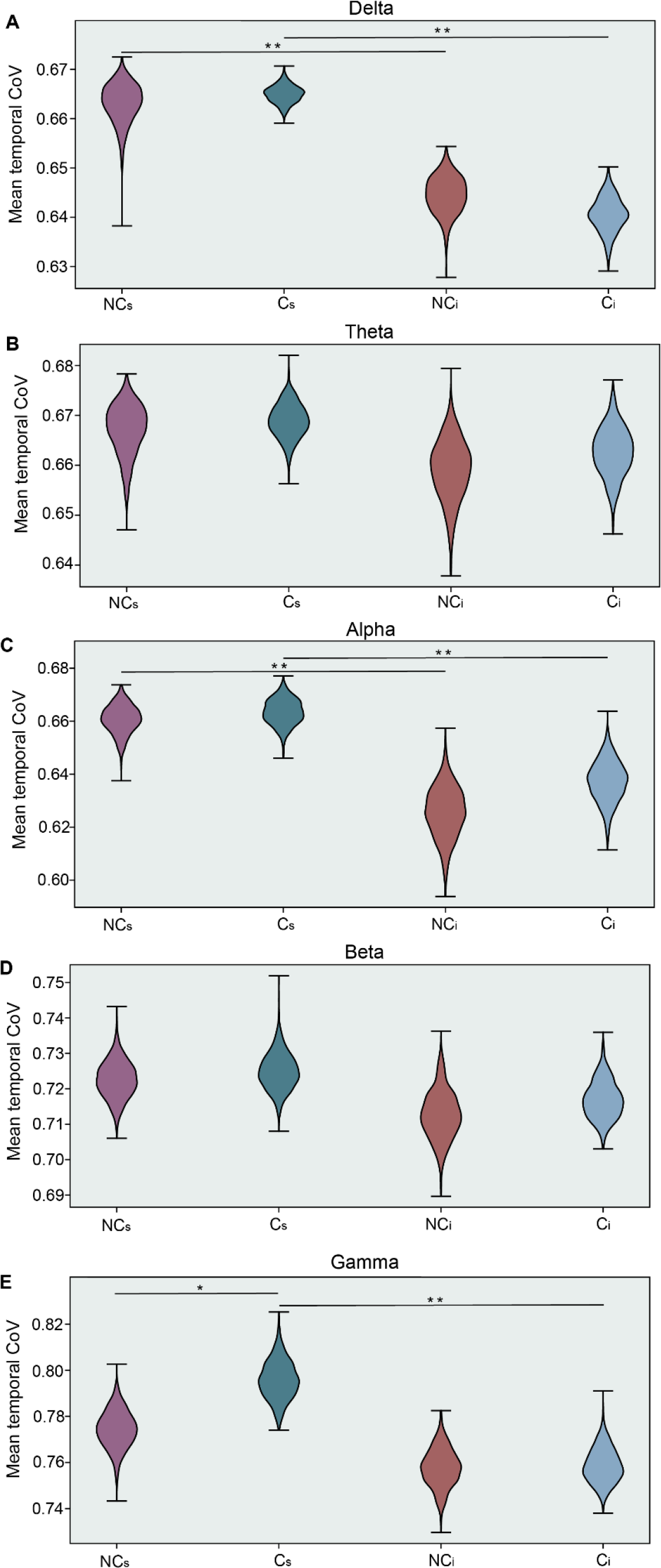
Social Media and in-person discussion comparison. (A-E) Mean wPLI temporal coefficient of variation (CoV) for each frequency band including (A) delta,(B) theta, (C) alpha, (D) beta, and (E) gamma. Each panel presents results for the intervals with no opinion change (NC: purple-red) and with a change in opinion (C: blue-green) interacting with the social media platform (C_s_,NC_s_) and during in-person discussion (C_i_,NC_i_). Statistical differences were estimated through a bootstrap procedure (1000 replicas with N=30 each) with significant results denoted with asterisks and are shown both within group and across social interaction contexts (* *p* < 0. 05, ** *p* < 0. 005).

### Summary of Findings

In aggregate, we have observed varying levels of sensitivity in the estimated metrics, attempting to describe the neural dynamics underlying social context and opinion change influences within the brain. Due to the complex nature of the findings, the differences in context and the cognitive processes driving our findings, we aggregate and visualize our results in Figure 5. We present our findings along 2 continuous axes and 1 static axis. First, we showed results for 3 different network metrics: *flexibility,* or the propensity of a node to change its affiliation across time, *allegiance*, or the pairwise likelihood that two nodes are in the same community, and *intermittence*, or the rate at which pair-wise affiliation change (1 = constant and consistent change, 0 = no change). These three metrics are ordered as a function of how spatio-temporally resolved they are. Flexibility, a node-wise metric, does not differentiate between pairwise similarity and/or differences in community affiliation. Allegiance, on the other hand, is a pairwise measurement of “similarity” but does not take into account how node-pairs change their affiliation across time. So, these three metrics are on an axis representing low spatio-temporal resolution at the origin and high temporal and spatial resolution at the top portion of the graph. As our results indicate, these metrics must be interpreted within the context of the specificity in frequency band, where each band has been associated with a variety of cognitive phenomena, but also represents communication across long distances (e.g., delta) or short (e.g., gamma). For the least spatially resolved metric, *flexibility*, the differences between the change and no-change groups was most robust within the beta band, a frequency band often implicated in motor behavior (e.g., Stancák & Pfurtscheller, 1996), but also in signaling the “status quo” (Engel & Fries, 2010). While allegiance, by itself, did not reveal significant pairwise changes (see Supplemental Figure S2), considering intermittence at varying levels of allegiance revealed the most robust changes across the groups at each frequency band of interest. Interestingly, intermittence, by itself, was most robust in the highest frequency bands. Together, these results indicate a specificity to these metrics but a robustness of the *intermittence* metric, suggesting a unique importance to the nature in which networks reconfigure in a complex decision making task, like opinion formation and/or change.

**Figure 5:**
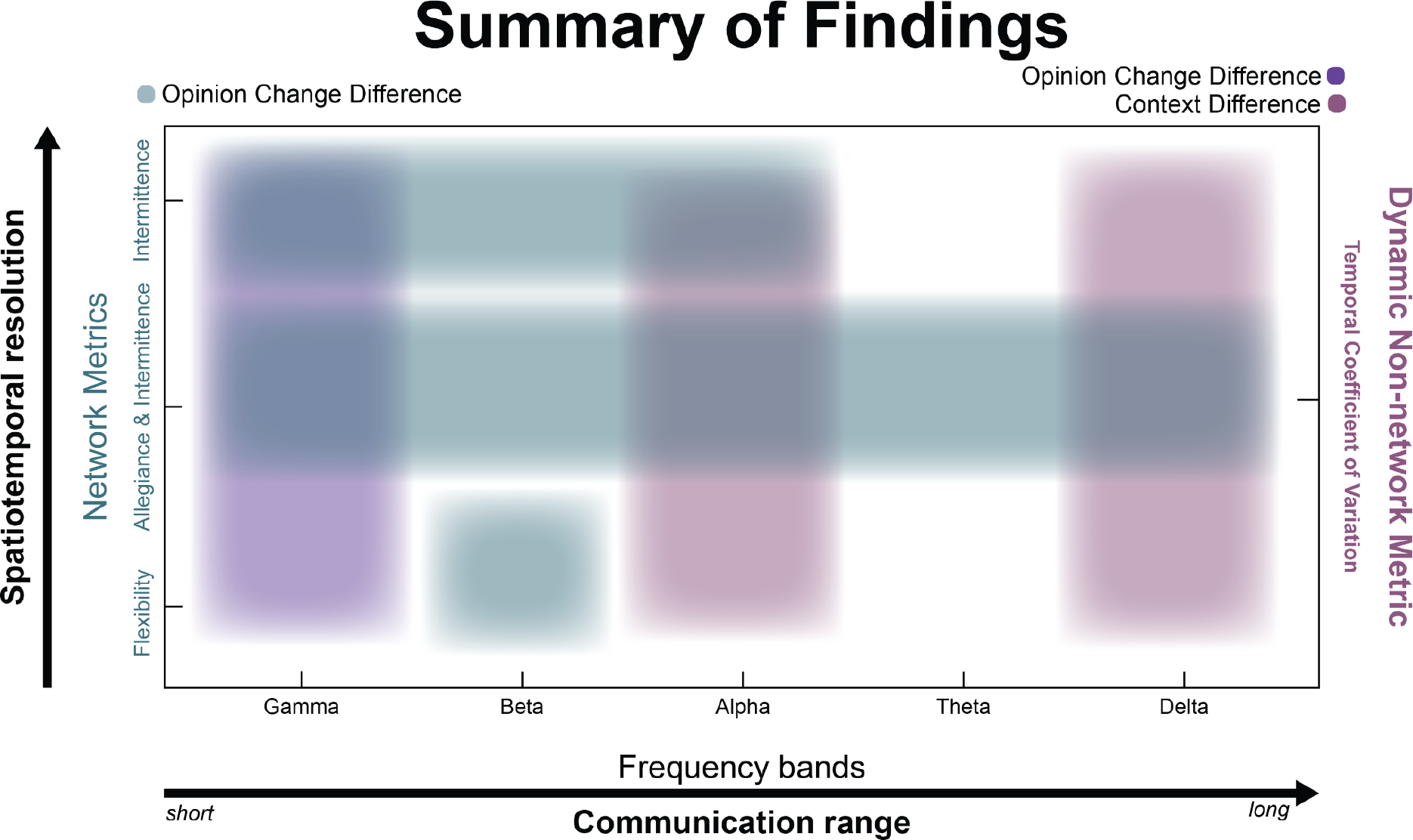
Summarizing the dynamics to opinion change. In aggregate, we summarize the findings along three axes; (i) one in which the network metrics describe the spatiotemporal resolution of the network dynamics (y-axis, left), (ii) an axis meant to describe the effects within the estimated frequency bands, interpreted as short to long range communication within the brain (x-axis, bottom), and finally, (iii) a proxy for flexibility that estimates the overall dynamics across the scalp as measured with coefficient of variation (y-axis, right). Network metrics on the left y-axis are sorted by increasing ability to resolve the nature of the spatial specificity and temporal dynamics. On the x-axis frequencies are sorted in descending order in terms of communication range where gamma is sensitive to changes in proximal neural ensembles and delta to long range communication. Purple hues should be associated with the right axis and teal hues are meant to be associated with the axis on the left.

As a final summary, we also inspected a proxy for these dynamic metrics, specifically inspecting the mean coefficient of variation (CoV) across time of the estimated functional connectivity (wPLI). Here, a high CoV is associated with a fluctuating connectivity and may be associated with how often a node changes it’s affiliation across time. Indeed, we see that this metric can capture the variance associated with changing one’s mind after the social media interaction, but only within the gamma band. Interestingly, this metric also captured variability in context (social media vs in-person discussion) within the alpha and delta bands, suggesting perhaps a different mechanism by which context shapes the way we encode information and make decisions.

## Discussion

Decision making is a complex internal process by which information is consumed and an action is executed, requiring the support of many interacting brain networks composed of a variety of functionally diverse regions within the brain (for review, see Fellows, 2004; Rilling and Sanfey, 2011; Wallace and Hofmann, 2021). The present study investigated the impact of informational context and its type on decision processes, specifically how social media and in-person discussion influences one’s malleability to *change one’s mind* on “highly shared” content in online platforms. Our findings have shown a large portion of individuals (e.g., 5 out of 6 in vaccination hesitancy) were susceptible to content displayed to them on a simulated social media platform while the same individuals were not susceptible to freeform and in-person discussions on the same topics. Using dynamic community detection (see Garcia et al., 2018 for a review) and estimating the temporal dynamics of network reconfigurations that occurred across several frequency bands, we found that the flexibility of specific sensors across the scalp could discriminate between those individuals who were and were not influenced by content presented in a simulated social media platform (Fig. 2). Importantly, those with no change in their opinions showed higher flexibility in sensors located within the frontal and posterior regions for the higher frequency bands (i.e., alpha, beta, gamma) whereas, the opposite effect is observed in theta band where higher flexibility was observed for those in which there was a change in opinion within prefrontal and posterior sensors (Fig. 2). Our results extended the flexibility findings to show that the vigor in which the network changes occurred was driving this effect with a new metric we call *intermittence*. Interestingly, intermittence within the higher frequency bands was more robustly diagnostic for different opinionators than lower frequency bands (Fig. 3). Moreover, while our results did not show substantial change in opinion after in-person discussion (only 35 people changed their mind after in-person discussion), we also did not find any of the observed brain network reconfiguration changes during the in-person interaction (Fig. 4).

### Network flexibility as a marker of opinion change

The so-called *flexibility* metric has been used to describe the rate of motor learning (Bassett et al., 2011; Bassett et al., 2013; Gerraty et al., 2018; Li et al., 2019; Reddy et al., 2018), has been associated with multitasking (Alavash et al., 2015; Shafiei et al., 2020; Thomas Yeo et al., 2015), pattern recognition (Telesford et al., 2016), language comprehension (Chai et al., 2016), thought rumination (Han et al., 2020; Lydon-Staley et al., 2019), adaptations to new stimuli or stress (Paban et al., 2019; Betzel et al., 2017), and working memory (Braun et al., 2015; Lauharatanahirun et al. 2020). Our findings support the increasing evidence suggesting the importance of the rapid reconfigurations of brain networks in cognition, and specifically, here, decision making as it pertains to opinion change and/or formation (for review, see Shine and Poldrack, 2018). We used *flexibility* as the probability a particular node (i.e. sensor) changes its affiliation across time. Previous studies show that this type of network-defined flexibility in frontal brain regions is associated with faster motor learning (Basset et al., 2011), psychological resilience (Paban et al., 2019), chronic behavior change in addiction (Cooper et. al., 2019), enhanced working memory performance (Braun et al., 2015) -- which is also moderated by sleep (see Lauharatinahirun et al., 2020) -- and even improved adaptive problem solving skills (Barbey et al., 2018). Given the diversity in these behavioral findings and our extension to even social media influence, it is reasonable to attempt to understand how this metric may be highly sensitive to a variety of complex cognitive phenomena.

Indeed, the neuroscientific journey that has led to the importance of *flexibility* in neural behavior may be understood from several different perspectives, and it is currently thought to be the foundation to the human’s unique ability to rapidly adapt to task demands. First, it should be noted that we have estimated network-based flexibility via a mathematically defined dynamic network approach (see *Methods*) and on its surface it should not be confused with concepts such as *cognitive flexibility* (Uddin, 2020) and *neural flexibility* (Yin & Kaiser, 2021), but it can be complementary to both (Mattar et al., 2016). Cognitive flexibility refers to the executive functions that allow an individual to rapidly transition from task to task and has been found to be associated with improved performance in a variety of tasks and also is reduced in certain pathologies (Hanes et. al., 1995). In contrast, neural flexibility, while often used in relation to cognitive flexibility, refers to the brain’s ability to rapidly shift across tasks and be recruited for a variety of activities (Uddin, 2020). Dynamics in the neural signal have previously been discarded as noise, but are now accepted as describing valuable variability in human behavior and even psychopathology (Uddin, 2020). Our findings not only add to this growing literature and support the network science approaches that can successfully capture this variability, but specifically, we have used dynamic community detection, an extension of the network science approach of functional *modularity,* that is a theoretically derived (but individualistic) technique to probe dynamic network changes via a distillation of dynamic connectivity matrices (Garcia et al., 2018). Our research adds to the general nature of this technique to capture the broadly cognitive, distributed and adaptive nature of the brain, the primary criteria for flexible brain regions and networks (Yin & Kaiser 2021). Here, we speculate that the process in which we are rigid in our opinions shows faster network reorganizations due to an effort to accommodate conflicting information with previously held beliefs. Another possible explanation can be different social media browsing strategies that led them to be exposed to more content for a smaller amount of time, being less prone to reevaluate previous beliefs, while increasing the sensory input. Future studies may be designed to evaluate these options.

### Intermittent and persistent network reconfigurations are diagnostic of opinion change

Our results expand on the pervasiveness of the flexibility results. We show that, at least for complex, high-level decision making, not only the rapidly evolving network reconfigurations (as measured by flexibility) are important but also the fast dynamics of intermittent linking (same nodes linking in an on-off fashion, a.k.a. *intermittence*) between modules are more diagnostic of social media influence on one’s opinion. Dynamic community detection has proved to be an effective tool to explore temporal patterns in systems represented by complex networks and a key aspect for this framework is the determination of the temporal resolution of the dynamic communities (Telesford et. al. 2016). A systematic way to determine a temporal resolution which leads to behaviorally relevant network description of the brain can be achieved by modularity maximization (Newman, M E J, 2006). Based on the dynamic communities obtained through the modularity maximization algorithm, we explore the temporal patterns of network links and how community allegiances of the network nodes change across time.

A critical feature of our findings is the fact that the temporal profiles of the estimated community structure is more diagnostic (e.g., *intermittence)* than simply the fact that dynamic network reconfigurations occur (Fig. 3). The temporal profile of interactions has a fundamental importance on a wide range of phenomena such as the dynamics of neuron populations that lead to seizures (Jirsa et. al. 2014, Rungratsameetaweemana et al, 2021), weather models and turbulent systems such as the Lorenz attractor (Ruelle 1976) and the many synchronization phenomena in which many units share the same temporal profile (Pikovsky et. al. 2001; Strogatz, 2000). From the point of view of dynamical systems, processes of opinion spreading have been extensively studied using models such as the Voter (Holley and Liggett, 1975) and Majority rule models (Krapivsky & Redner, 2003), suggesting a complex interactive scheme that gives rise to opinion formation and change. Interestingly, recent findings suggest that information sharing and spreading occurs at a faster pace in social media platforms than in-person social contacts and explores the effects of these two time scales in a consensus formation model (Ding et. al., 2018). With our approach, we explore this opinion change phenomenon at an individual level using the complex networks framework to identify connectivity patterns of EEG data that are diagnostic to an opinion change process during a social media interaction. In aggregate, these findings coupled with our current results, suggest similar operations at both the neural level and population level. Recent findings suggest the brain (as well as other complex systems) operate outside of the boundaries of a particular spatial scale (Cocchi et. al., 2017). Perhaps, our results are the consequence of information spread, whether within a single brain or across human interactions and are suggestive of a scale-free phenomenon (Mahmoodi et. al., 2017). Indeed, there are many complex systems that express this scale-free behavior; however, it should be noted that recent findings have even shown that this universal principle is flexible (Bansal et. al. 2021).

### Network reconfigurations and oscillatory specificity suggest a complex operation within the brain in opinion change

Despite the increasing efforts to understand the neural processes that underlie deliberation and decision making, much on this subject remains unclear; however, important findings from the literature in EEG oscillations, evidence accumulation, valuation and identity may play critical roles in understanding our results. First, we have shown that intermittence effects are more robust at higher frequencies than lower frequencies in the observable EEG oscillatory scheme (e.g., gamma vs theta). We also show that the delta band is most diagnostic for social interaction context. Oscillations emanating from the brain, as measured with EEG, are a consequence of short- and long-range connections within the brain that interact to give rise to cognitive capabilities (Buzsáki 2006). Importantly, the slower oscillations mostly represent the coordination of distal regions within cortex and sometimes even modulate higher frequency oscillations within the brain (e.g., Bragin et al., 1995; Chrobak and Buzsaki, 1998; Leopold et al., 2003; Schroeder and Lakatos, 2009; Canolty et al., 2006; Buzsáki and Wang, 2012; more recent ones). In other words, oscillatory activity and the associated cognitive functions rely on the global coordination of local processes (Cavanaugh & Frank, 2014; Rungratsameetaweemana et al., 2018). Within this context, it would suggest that our results could be a consequence of both, where the most critical “intermittence” effects were observed within the delta and gamma bands, “flexibility” effects were most critical in the beta band, and social interaction context was most observed within the delta band.

This broad coordination of neural communication in the brain gives rise to specific cognitive functions, and our results could reflect several different operations at play. First, our findings show the most significant results in flexibility within the beta band. The beta band is often associated with motor behavior (Khanna & Carmena, 2015), but has also been proposed to carry a more prominent role in maintaining motor or cognitive states (Engel & Fries, 2010); interestingly, beta band dynamics have even been associated with the accumulation of evidence in the sensorimotor network in a vibrotactile decision task (Haegens et. al. 2011). Moreover, fMRI and transcranial magnetic stimulation (TMS) studies have extensively implicated the so-called *value system* in decision making, a system that is engaged when weighing the potential benefits of a particular decision route. Critical components of this system are thought to include the ventromedial prefrontal cortex (VMPFC) and ventral striatum (VS). These regions, within social contexts, have been linked to susceptibility to social influence from peers (for a comprehensive review see Falk & Scholz, 2018). More broadly in EEG, several frequency bands have been implicated in decision making (e.g., Nakao et. al. 2019), but often show some specificity in frontal and parietal regions (e.g., Golnar-Nik et. al. 2019). Interestingly, a recent study inspecting long range temporal correlations (LRTC) in EEG recordings has shown a relationship between theta to alpha bands and the abstract concept of self identity and identity confusion (Sugimura et. al. 2021). Due to the highly complex decision, speculation, and potential action, our results most likely indicate a complex coordination and reconfiguration of networks within the brain, across several frequency bands and reflect coordination of these processes including evidence accumulation, valuation and even internal reflection on identity. Future research is needed to disentangle these processes and influences on decision making context, especially within the social media and in-person social contexts studied here.

### General properties of network reconfigurations within the brain

A somewhat unintuitive result is the finding that those individuals with more rapid network reconfigurations are associated with a lack of change in opinion. We previously speculated that this may be due to an effort to accommodate conflicting information with previously held beliefs or mark different social media browsing strategies that led them to be exposed to more content for a smaller amount of time. We hope that future research can differentiate between these potential mechanisms, but when considering our findings within the ubiquitous nature of these network reconfigurations and associations with behavior, we believe it may be a more generalized process than previously understood. For example, let us consider a recent theory of general intelligence based on how we navigate spatially through the world (Hawkins et al., 2019). This theory is consistent with several principles that include generalized machinery of the brain (Mountcastle, V., 1978) to navigate throughout whatever real (or abstract) space, prepare predictions (Rao & Ballard, 1999), and compare to some reference frame built from previous experiences (Lewis et al., 2019). It could be the case that metrics derived to estimate the dynamic reconfigurations within the brain are targeting this rapid navigation through possible “observational” (sensorial) interpretations and possible actions, a foundational element to not only critically think about a topic but also is fundamental to general intelligence. Given that, we hypothesize that those that are highly flexible are also navigating through this abstract sensory, perception, decision, and/or action space more rigorously than those that are not.

### Methodological considerations and future directions

Within this manuscript, we prioritized the ecological validity of the experimentation rather than distinct cognitive constructs. While the latter presents a very interesting avenue of scientific pursuit, introducing controlled conditions or other contrived experimental manipulations could potentially modify behavior in unexpected ways. Indeed, often with ecologically valid research, there is a limitation in understanding all elements that have contributed to our findings. For example, each subject was able to navigate freely through the social media content, and there was no control on how much time was spent with the content, so variable input may be a potential confound; however, traditional laboratory-based experiments are not only defined by their highly precise neural findings, they could also shape the responses in unnaturalistic ways. Similar to the trends in social (and general) neuroscience studies (Osborne-Crowley, 2020; Pawel et. al. 2019), we believe that our results should be complimented by more laboratory-specific effects. In other words, our results don’t take into account any content-specific processes or differences on individual social media interaction, nor do they consider the controversial and emotional aspects of the different scenarios presented for the subjects.

Within this experiment and the design that prioritized ecological validity, there are also several considerations in the experimental measurements (e.g. EEG) and the specific type of opinion change that was observed of note for future experimentation. For example, the type of EEG connectivity that was deployed in this analysis, prioritized the phase-based statistical dependencies (for an additional related results, see Figure S5 in Supplemental Materials) within *a priori* and somewhat narrowly defined bands of interest, following traditional techniques and to compare to the wide literature in oscillatory action as measured with EEG. Moreover, as is common with EEG analyses, there is a potential that multiple sources may contribute to the effects, as the spatial information from EEG is known to suffer from volume conduction. The high temporal resolution of EEG, however, creates an opportunity to extend dynamic network measurements to more rapidly evolving network reconfigurations (as compared with fMRI) and in a less constrained manner (e.g., at a desk, in a group discussion rather than scanner), the inherent limitations in the spatial resolution and the potential contributions to the signal should be considered in future studies. While the combination of flexibility, intermittence, and coefficient of variation analyses converge to a clear importance of the dynamic network reconfigurations within the brain and their relationship with opinion change, different modules within the brain may be sensitive to different cognitive phenomena. Future research may extend these findings to other cognitive actions that contribute to this complex decision, perhaps even further disentangling the sub-elements of this process including perception, weighing alternatives, belief consistency, social influence, etc. Moreover, within our study, the overwhelming contribution to the type of opinion change involves determining the consequences of a murder trial, of which, little information is directly known to the subject and created a wide range of responses (see also Figure S4 in Supplemental Materials). The generalizability of these findings is yet to be known; perhaps, even, the non-personal nature of these results have a substantial impact on the findings. This might be an aspect of the findings to be assessed with further experimentation.

Within this context, our results are merely the first step toward understanding the dynamic reconfigurations within the brain and how different context and content interact to give rise to opinion change, highlighting the difference between in-person discussion and social media interaction. Future studies will investigate the unique aspects of opinion change that are generalizable beyond the scenarios presented here, include within-subject condition comparisons to inspect general properties of opinion change within these two social contexts, and perhaps even understand the interaction of human biases in their interaction. For example, self-referential opinion changes may suffer from the interesting optimism bias (e.g., Sharot, 2011), thus requiring different neural resources than non-self-referential.

Other methodological limitations are related not to experimental design but to the analytical technique in which we estimated the dynamic network reconfigurations. Here, we use dynamic community detection (for review, Garcia et al., 2018) to distill connectivity patterns derived from phase-based statistical dependencies into communities, or clusters of electrodes and then estimated shifts in communities across time. We have previously extended this method to EEG on a limited number of channels (Garcia et al., 2020), but there are lingering questions on robustness of the method to varying number of channels, the cognitive aspects the chosen temporal window might capture, and the parameter search. Future studies may explore other techniques to capture the processes underlying opinion formation, change, and generally complex decision making. Moreover, future studies are needed to understand full contributions to session and subject level variability as well as disentangle the “intermittence” results as potentially marking different internal processes while interacting with social media or different strategies deployed for social media interaction.

## Conclusion

The current study used a complex network based framework (dynamic community detection) to investigate the relationship of brain dynamics during social media interaction with the opinion change and/or formation processes. Our results indicate that the rapidly evolving network dynamics in delta, beta and gamma bands are the markers of influence of social media platform interaction on opinions in a range of scenarios, such that the slower dynamics is associated with individuals who are more likely to change their opinion. We also introduced a new metric called *intermittence* to assess differences in the observed faster or slower network dynamics. Estimating the intermittent and persistent network changes (as measured via *intermittence),* our results suggest that the functional brain network structure for opinionators with opinion change also show differences when interacting with social media platforms compared to in-person discussions. Together, our results suggest unique decision making operations during social media interaction and represent trait-like dynamics in individuals that change or not in their opinions.

## Methods

### Participants

The data was collected from a cohort of 123 healthy participants between the ages of 18-40 years. Subjects were screened and the ones diagnosed with sleep, psychiatric, neurological, eating, behavioral (including Attention Deficit Disorder), or cardio-pulmonary/vascular disorders, uncontrolled blood pressure, heart disease, HIV+/AIDS, head trauma within the past 5 years, regular use of prescription drugs that can alter EEG or impair their ability to participate, use of illegal drugs (recreational and medical marijuana users were not excluded), excessive use of nicotine, alcohol and/or caffeine, untreated vision or hearing issues, pregnant or nursing, and inadequate familiarity with the English language were not included. For more information on the recruiting procedure we refer the reader to (Richard et. al., 2021). The data acquisition for this study using human participants was reviewed and approved by Alpha IRB and Air Force Research Laboratory (AFRL). The participants provided their written informed consent to participate in this study.

### The Social Media Platform

This study used an innovative platform known as the Social Media Analytic Replication Toolkit (SMART). SMART, which was developed by The Intific Division of Cubic Defense Applications Inc., (CDAI), and allowed users to interact within a closed social media environment for experimentation and real-time exercises that was inspired by Facebook and Twitter. Social media accounts were created for each of the group participants which allowed the participant to like, share, or retweet posts as they normally would do on their personal social media accounts. Participant interactions with SMART were not broadcasted to other group participants’ accounts, but all other aspects of the platform were meant to make the “feel” of the platform as realistic as possible. For screen shots of the platform, please see Supplemental Figure S7.

### Procedures

For the experimental timeline and descriptive analysis of behavioral results (e.g., opinion change), see Figure 1. Briefly, after arriving in the laboratory, subjects were presented with questionnaires that asked their opinions on particular real-world scenarios found commonly on social media platforms. Questionnaires were presented before and after interaction with the simulated social media platform as well as after the in-person discussion segment. EEG was recorded during these two interactive contexts, i.e., *social media interaction* and *in-person discussion*. To briefly describe the procedures, participants were seated in front of a computer after instrumented with EEG and allowed to freely browse through the 3 scenarios and interact (e.g., like, share, etc) with the content freely. While the platform contained 4-5 articles that could either skew the opinion of the individual in two different directions (e.g., to vaccinate or to not vaccinate), there were no restrictions or trials; however, the social media interaction was limited to 2 hours and the in-person discussion segment was limited to 20 minutes. The in-person discussion was administered directly following the completion of the second questionnaire and participants were directed to a room to discuss each scenario. A research technician was present and acted as a moderator to ensure that the discussion was efficient and appropriate and all topics were discussed for approximately 5 minutes. Each scenario was addressed in the same order for all groups during the discussion period. Participants were asked the same two questions for each scenario by the moderator: “What did you decide and why?” and “What, if any, social media posts influenced your decision?”. When the discussion ended, the participants were instructed to complete scenario-specific questionnaires one last time before completing their participation in the study.

### Scenarios

Three hypothetical scenarios were presented to the subjects. During each session, subjects were exposed to the contents of three scenarios. The opinions of each subject in the scenarios were accessed through questionnaires that were delivered before and after interaction with a social media simulation software. The following details the three scenarios analyzed in this study.

#### Free travel destinations

Within this scenario, subjects could choose between two locations (Paris, France, or Sulawesi Island, Indonesia) for an all-paid one-week vacation where each location was vulnerable to different dangers; Paris could have large protests and sporadic violence and Sulawesi Island had the potential for a destructive tsunami. Articles presented to the subjects mentioned these dangers and mentioned how nice the locations were to visit, with equal representation. The subjects were also given the opportunity to volunteer, in support of the rebuilding effort following the catastrophic damage; they were also asked how much time they were willing to dedicate to the rebuilding effort.

#### Murder trial

In this scenario, subjects were asked to imagine themselves in the jury of a trial, which was based on a true story. After receiving information about a case in which a young female college student was murdered, they were asked whether the young man accused of murder should be considered guilty or not, and if guilty, the length of the sentence or death.

#### Vaccinations

In this scenario the subjects were asked about vaccinating one of two hypothetical children after the older one started to show development impairments after being vaccinated. The questionnaire presented only the binary question if the subject would vaccinate or not their second child, and the online content was equally for or against vaccinations.

### Behavioral metrics

The subjects answered questions about each of the hypothetical scenarios and opinion changes on the topics of the three scenarios were evaluated to determine the likelihood of opinion change. Of the three scenarios, the *vaccination* scenario was the only completely binary response as the *travel and murder* scenarios included followup questions that were not binary. Thus, several steps were completed to include all three scenarios in the analysis and construct a behavioral metric of opinion change.

For the three scenarios analyzed in this study, the social media opinion change was coded as either a 0 (no change) or a 1 (change), indicating a change in opinion relative to the previously answered questionnaire. In other words, for the social media interaction, change was measured relative to the first questionnaire and for the in-person discussion condition, opinion change was assessed relative to the second questionnaire (completed after social media interaction). Change was coded as follows: for the *travel* scenario, a response was considered a change (1) in opinion if any of the following were true: (i) the subject changed the destination choice from France to Indonesia or vice versa or (ii) the subject modified their decision to volunteer. For the *murder* scenario, a response was considered a change if any of the following were true, (i) the subject changed their opinion from guilty to not guilty and vice versa, (ii) the subject changed their opinion on prison time or punishment. For the *vaccination* scenario, the responses were either yes or no, so a response was considered a change if it did not match the previous response. If the answers are all identical to the previous survey, then the response was coded as a 0, or no change. The social media opinion change score for each subject was estimated as the sum of the social media opinion change scores of each individual scenario and could be 0,1,2, or 3, where a 3 represents a change in every scenario and 0 in none. Critically, groups were defined as those that did not change their opinion in any scenario (no change) and those that changed their opinion in at least 1 scenario.

### EEG Analysis

#### Preprocessing

EEG was acquired using the B-Alert R X24 wireless sensor headset (Advanced Brain Monitoring, Inc., Carlsbad, CA, United States) placed on the subjects before the subjects interacted with the social media platform. The headset is composed of 20 EEG sensors located according to the International 10–20 montage at Fz, Fp1, Fp2, F3, F4, F7, F8, Cz, C3, C4, T3, T4, T5, T6, O1, O2, Pz, POz, P3, and P4. Reference sensors were linked and located behind each ear along the mastoid bone region. The sampling rate was 256 Hz and the signal was filtered through a high band pass filter at 0.1 Hz and a low band bass, fifth order filter, at 100 Hz. To insure high quality data was collected, a maximum allowable impedance was set to 40 k . Next, the data was band-pass filtered within common frequency bands including delta (1-3 Hz), theta (3-7 Hz), alpha (8-13 Hz), beta (21-30) and gamma (25-40) using a low order (3) zero-phase forward and reverse digital IIR filter in Matlab (Mathworks, Inc.).

#### Temporal Windowing

Temporal windowing is a common procedure in studies of EEG, where it may capture (1) the temporal scale of a variety of phenomena, (2) may indicate the specificity and robustness of the chosen temporal window (10s) in relation to the cognitive phenomenon in question, and (3) lies at the intersection of measurement sensitivity, environmental noise, and the covert mental events we study.

First, when considering functional connectivity, EEG has been shown to be sensitive to a variety of task effects (e.g., Nidal & Malik, 2014) but it is also well established that its feature capture more rapid changes in the brain (Garcia et al., 2020) and even more stable effects as may be understood with disease models and genetics (e.g., Smit et al., 2008), spanning a large variety of temporal scales and a variety of behavior.

Second, in relation to the chosen time window (10s), there’s a large literature that has sought to understand how temporal windowing choice can affect the ability to capture differences in relation to trait-based information (or individual differences) as measured with EEG. For example, previous studies have suggested that graph metrics, as estimated via connectivity patterns derived from EEG, are susceptible to bias in the actual value, showing for example, shorter path lengths for shorter windows (see Fraschini et. al. 2016, compare 2s vs 10s) but more stability above 10s window sizes. More critical to the analyses within this manuscript, however, is the relative nature of our connectivity matrices as they inform the dynamic communities that are estimated across time. In comparing the connectivity matrices of the individuals associated with each behavioral outcome (i.e., change, no change), there is no reason to believe differential noise or bias in the estimated connectivity as we are comparing them rather than interpreting, say, the value of the metric. Moreover, some research has shown an increase in robustness to noise of connectivity above 5s or so (see Bonita et. al. 2014). Importantly, as well, is that we use the genLouvain algorithm to estimate communities which has been shown to be more robust to noise, number of clusters, and number of layers than other algorithms (Bonita et. al. 2014). Thus, here we have sampled the functional connectivity over a robust time period with robust methodology.

Despite the evidence supporting our windowed approach, our general philosophy lies in a narrow interpretation of our results that accounts for the nuances of our measurement technique, our methodology, the environment (or context), and the phenomenon in question. For example, often the aforementioned robustness studies are either simulated or constructed from resting state data, often trying to understand the stability of these metrics within an individual or across a population. While this approach is sound, it should really only be considered within the context of the resting state events, which are associated with mentalizing (Hein & Singer, 2008). Critically, as well, we interpret our results with specific frequency bands, where we assume low frequency band phenomena to be sensitive to long range communication within the brain and high frequency band effects are capturing localized and rapid communication within the brain. This has large impacts when interpreting the window size and the frequency bands that show these effects. For an extended analysis of different window size, please see Figure S3 in Supplemental. Importantly, the dynamics in connectivity, overall, can differentiate those that change and do not change their opinions within the gamma band at windows at 10s and above.

#### Functional Connectivity Analysis

To estimate the functional connectivity of the EEG recordings we calculated the pairwise weighted Phase Lag Index (wPLI) within each frequency band of interest, which is known to be highly sensitive to linear and nonlinear interactions (Imperatori et al. 2019). For each sensor, the EEG (already band-wise filtered) was partitioned in *L* windows with duration 10s. The dynamic changes of this 10 second window highlight differences between the two intervals in which subjects did and did not change their opinion. As has been observed Each time window was used to calculate a matrix in which each entry *A*_*ij*_ accounts for the weighted Phase Lag Index (wPLI) (Vinck et al., 2011) for the pair of sensors *i* and *j*, calculated as:

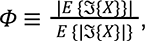

where *E* {.} denotes the expected value and 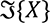 is the imaginary part of the cross-spectrum of the EEG recordings of sensors *i* and *j*. The temporal layers obtained by the described procedure were then used for the dynamic community detection analysis described in the next section. Importantly, though, the number of windows (*L*) were variable across subjects, with a mean *L* across subjects of 372 (SD = 100).

#### Community Detection and Network Dynamics Metrics

While human brain mapping efforts have demonstrated a relationship between spatial specificity and cognitive functions, techniques rooted in network science provide a useful framework for characterizing and understanding the spatiotemporal dynamics of the functional systems subserving cognition (Bassett & Sporns, 2017). One of the core concepts at the basis of network science is network modularity, which is the idea that neural units are structurally or functionally connected forming modules or clusters (Garcia et al., 2018). This organization allows for the system to perform both local-level exchanges of information, while maintaining system-level performance. Here, we examine whether a particular node’s propensity to change communities (i.e., flexibility) was related to change in opinions after interaction with a social network platform. To measure such changes in network communities during the interaction with the social media platform, a multilayer community detection analysis was employed (Bassett et al., 2011; Mucha et al., 2010) on the aforementioned wPLI estimates calculated for 10 s non overlapping windows (see Temporal Windows section above), with each social media interaction session accounting for 372±100 temporal layers. This method uses a Louvain algorithm to maximize modularity (Blondel et al., 2008) to define functional communities and is completed in several steps. First, it relies on two parameters, *ɣ* and ⍵, so called structural and temporal parameters of the analysis. We swept the parameter space from .5-4 for each parameter, subject, and segment and compared the mean estimated modularity value *Q* to a shuffled null dataset. We chose a parameter set that on average produced more than 1 community and was the highest difference in modularity from the estimated modularity from the shuffled null dataset (see Garcia et al., 2020; Garcia et al., 2020 for a similar procedure). This resulted in *ɣ = 1.1364* and ⍵ *= 0.5.* Due to the non-deterministic nature of the analysis, the chosen optimization procedure was repeated 100 times, since the algorithm is susceptible to multiple solutions (Good et al., 2010). From these multiple iterations, the following community metrics were computed: (i) *flexibility*, or proportion of time during which each node switches to a different community assignment; (ii) *allegiance*, related to how long two nodes are connected to each other during the task, and a new proposed metric (iii) *intermittence*, defined as how rapidly two nodes connect and disconnect through communities. Those metrics were calculated for each of the 100 iterations, and our results used the mean value for all the iterations. In more concrete terms, the flexibility of each node corresponds to the number of instances in which a node changes community affiliation, *g*, normalized by the total possible number of changes that could occur across the layers *L*. In other words, the flexibility of a single node *i*, ξ_*i*_ may be estimated by

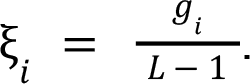

Allegiance is a metric calculated for each pair of nodes and accounts for the proportion of the total time a pair of nodes belongs to the same community, and is defined as:

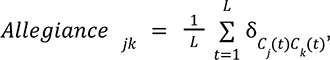

where δ denotes the Kronecker delta and *C*_*i*_ (*t*) denotes the community which contains the node *l* at time *t*. Therefore, 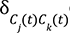 equals 1 if the nodes *j* and *k* are in the same community at time layer *t* and equals 0 otherwise.

Further, to account for the temporal dynamics of allegiance, we proposed a new metric, *intermittence,* which tracks how frequently the two nodes change their affiliation from the same to different and vice versa. Intermittence is defined as:

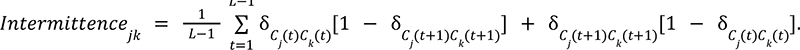

To visualize the concept of intermittence consider the example in Figure 5. First observe that the allegiance between nodes 1 and 4 is equal to the allegiance of nodes 2 and 3, however the link between nodes 1 and 4 is present for two large continuous epochs while the link between nodes 2 and 3 is connected for many short epochs, this characterizes the intermittence between nodes 2 and 3 as larger than the intermittence between nodes 1 and 4. Consider now the nodes 5 and 6, both have the same allegiance with node 1, however since node 6 changes its community assignment more often, its flexibility is higher than the flexibility of node 5. Observe that unlike intermittence, flexibility is a property of the node and is not calculated for individual links of the nodes.

**Figure 5:**
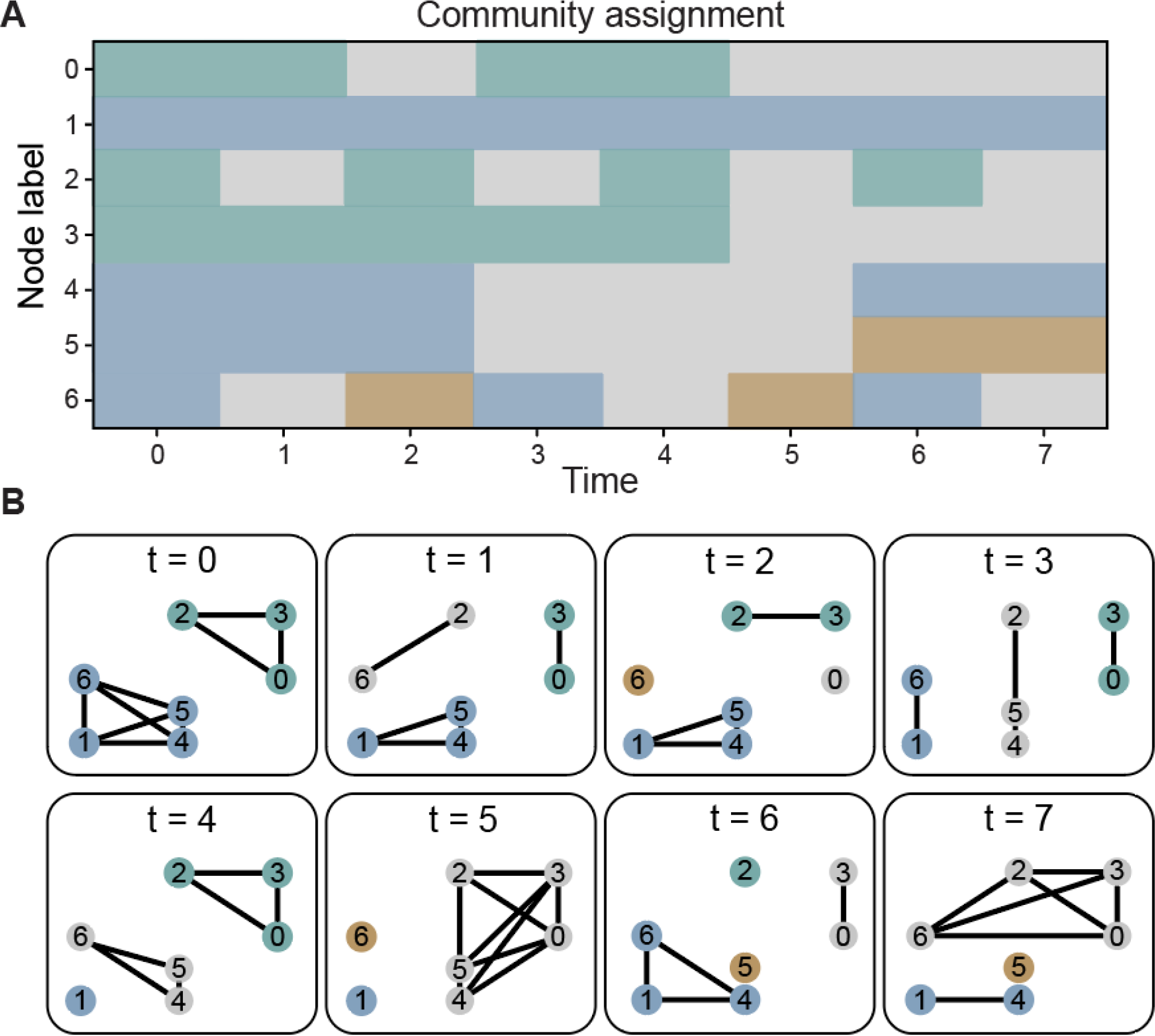
Flexibility, Allegiance, and Intermittence. (A) Example of a community structure assignment for 8 time layers. (B) Representation for each time layer of the community structure on (A).

#### Statistical comparisons

Two types of statistical comparisons were completed within the manuscript. Primary comparisons were between two groups, of unequal sizes, individuals who did and did not change their opinions. Due to the unequal sizes, bootstrap distributions (Wehrens et. al., 2000) were estimated and used to estimate p-value and 95% confidence intervals (Figures 2 and 3). For this method, 10,000 drawings (with replacement) were made within each group (change and no change) for each node (Figure 2) or across nodes (Figure 3) and means for each 10,000 replicas were calculated resulting in a distributions of the mean value (e.g., flexibility in Figure 2) or distributions of differences between groups (e.g., Figure 3B) were estimated. This process generated bootstrap distributions, from which 95% confidence intervals were then estimated. For the analysis in Figure 4, we carried out the bootstrap procedure with the random sorting on the individual level, calculating the CoV for each of the 1000 replicas with 30 randomly selected individuals each, and estimated the probability of the differences observed between the two groups for each condition, and for each group in the two conditions.

## Data availability

The raw data supporting the conclusions of this article and code will be made available by the authors upon request, without undue reservation. Neural metrics and behavioral change and scripts to reproduce the figures and analyses may be found in the Supplemental Materials.

## Acknowledgements

This research was sponsored by the US DEVCOM Army Research Laboratory and was completed under Cooperative Agreement Numbers W911NF2020067 (I.L.D.P.), and W911NF-17-2-0158 (K.B.). The views and conclusions contained in this document are those of the authors and should not be interpreted as representing the official policies, either expressed or implied, of the US DEVCOM Army Research Laboratory or the U.S. Government. The U.S. Government is authorized to reproduce and distribute reprints for Government purposes notwithstanding any copyright notation herein.

## Supplemental Material

### EEG sensor position

The EEG recordings were obtained using the B-Alert R X24 wireless sensor headset (Advanced Brain Monitoring, Inc., Carlsbad, CA, United States), the system has 20 channels and the montage layout is presented in (Figure S1). The reference sensors were located behind each ear on the mastoid bone. The sample rate was 256 Hz with a high band pass at 0.1 Hz and a low band bass at 100 Hz.

**Figure S1:**
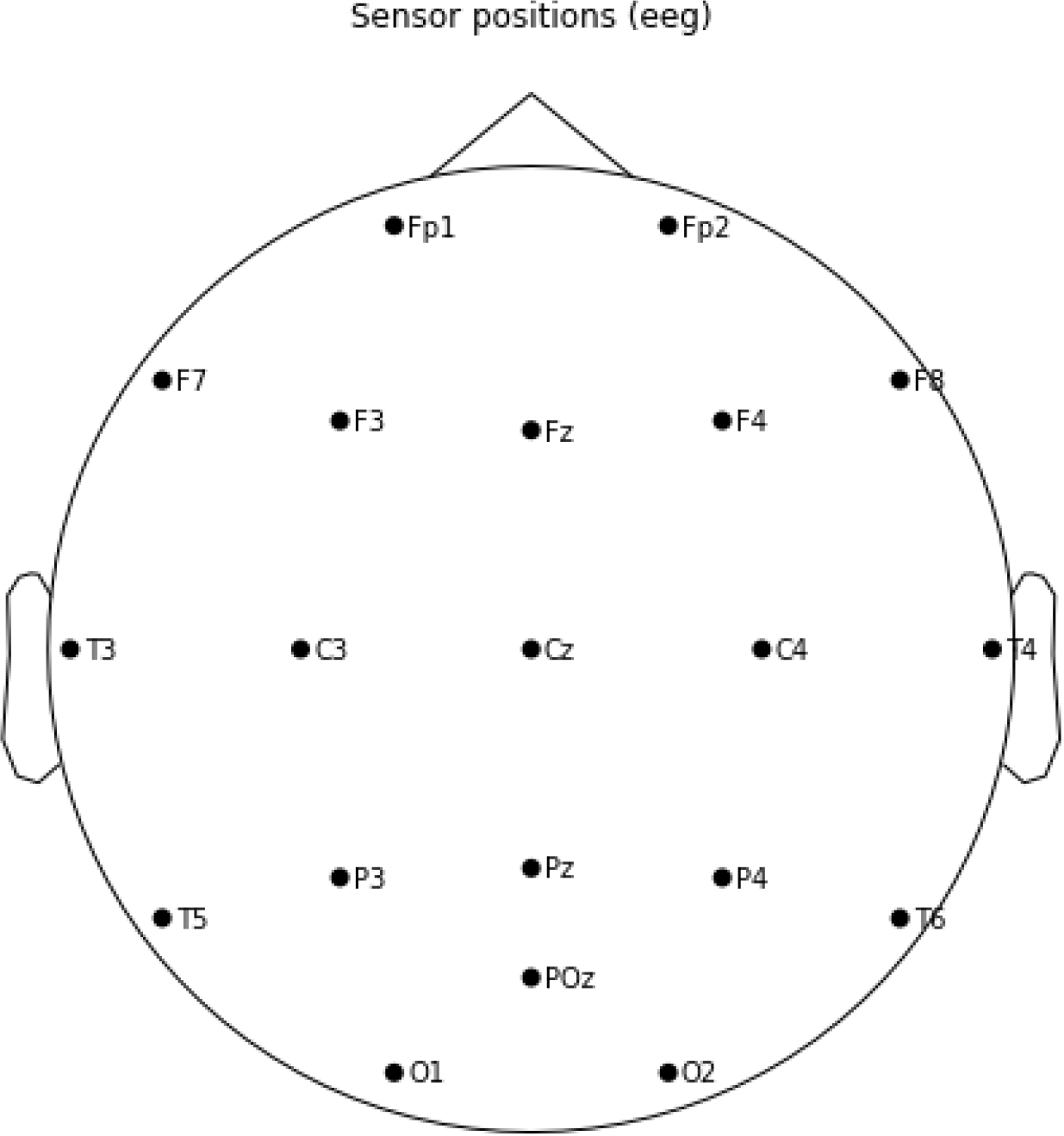
EEG sensor position. The topo plot shows the sensor montage. The EEG recording system model used was B-Alert R X24 wireless sensor headset (Advanced Brain Monitoring, Inc., Carlsbad, CA, United States) with 20 channels.

### Node allegiance

To further understand the differences between those that did and did not change their opinion in terms of network reconfigurations, we explored how the network nodes, particularly the nodes that showed significant difference in flexibility between two groups, (significant nodes, Figure 2) changed their functional allegiances. Allegiance is defined as the fraction of the total time two nodes are in the same community. First, we calculated the allegiance metric between all the node pairs and then compared them between two groups. None of the node pairs (including or excluding the significant nodes) showed a significant difference between groups. Moreover, on average, some of the node-pairs showed higher allegiances for those without a change in opinion while some showed higher allegiances for those who changed their opinion. In Figure S2, we show average allegiance differences between two groups. Yellow entries in the matrices represent higher allegiances for those with no change. In topographical plots we show these differences only for the significant nodes.

**Figure S2:**
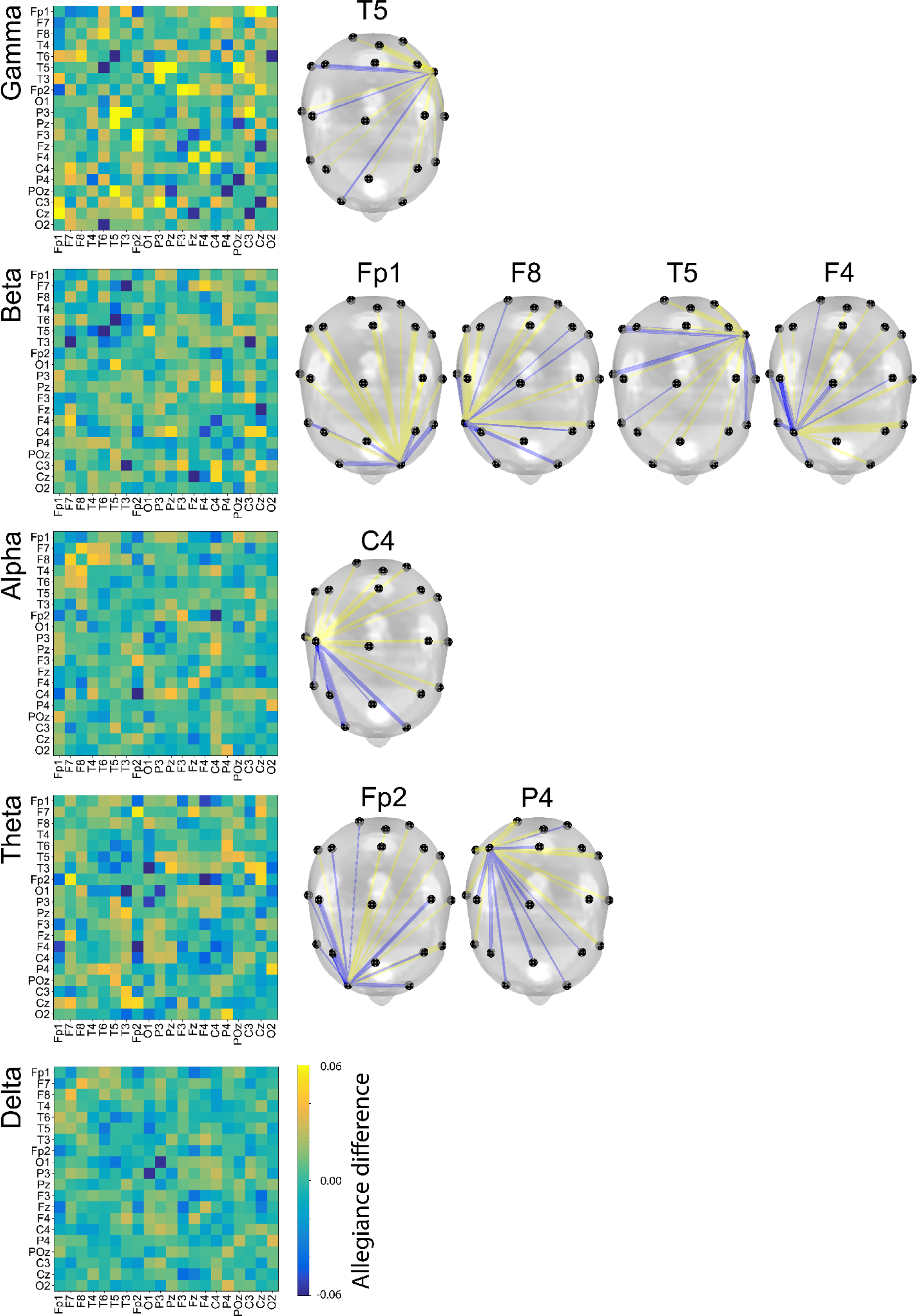
Node allegiance differences. The matrices show average allegiance differences between the two groups (no change and change). The topographic plots show the mean allegiance differences for sensors which showed significant flexibility changes between the two groups (as discussed in Figure 2). The links in yellow (blue) indicate a higher (lower) allegiance value for the no change(change) group.

### Effects of temporal window sizes

To explore the effects of different temporal window sizes we compared the mean wPLI temporal coefficient of variation (CoV) for the opinionators with and without opinion change during the social media platform interaction, the results are summarized in Fig. S3. As a general effect of the increase in the window size we observe higher values and broader distributions of the mean temporal CoV. The differences between the two groups captured on the gamma band are present for temporal windows larger than 10s, indicating that smaller temporal windows do not capture the temporal scale of the dynamics that reflect the different processes occurring in the two groups.

**Fig S3:**
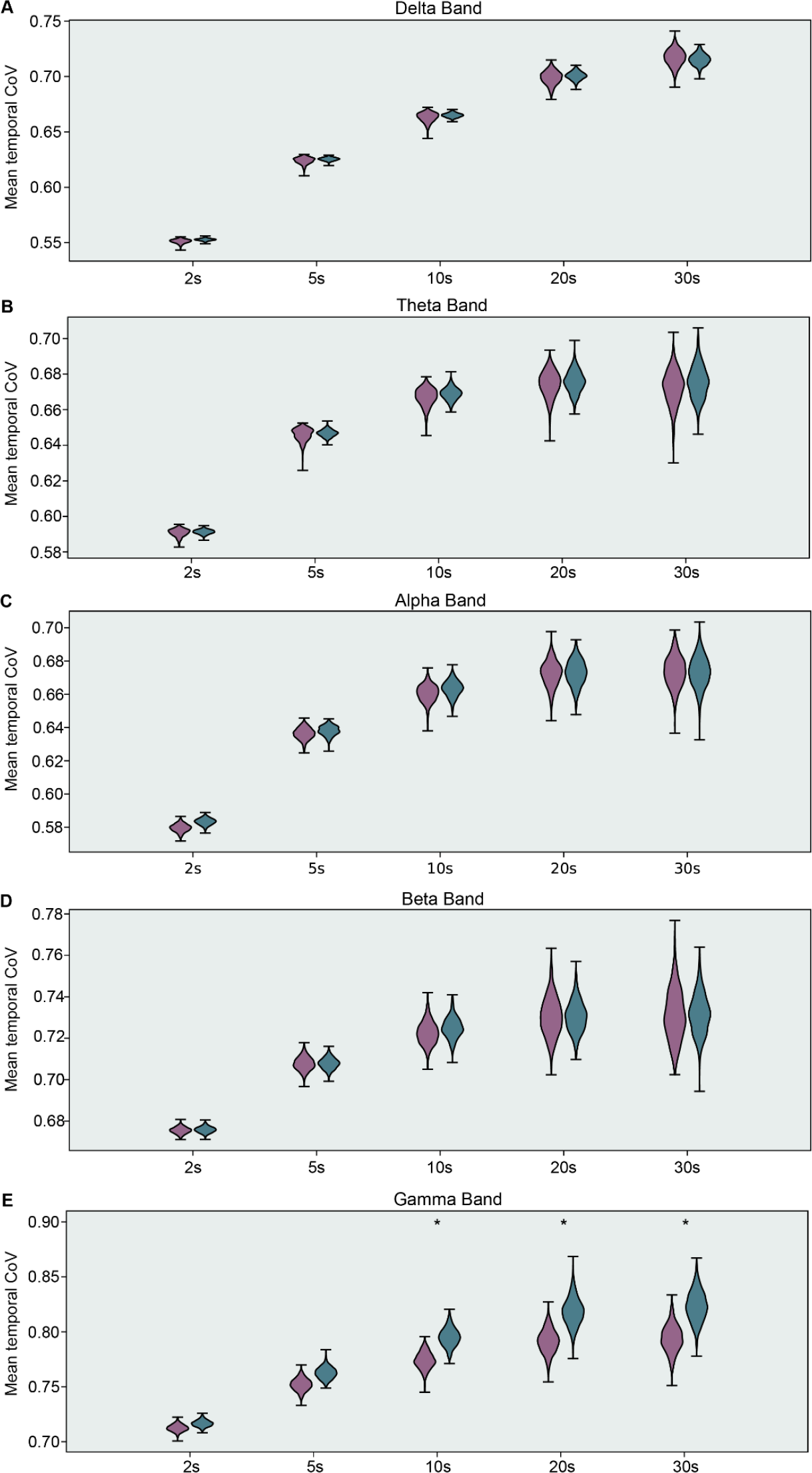
Effects of temporal window sizes. (A-E) Mean wPLI temporal CV for the those who do not change (purple) opinions and those who do (green) calculated for different temporal windows (2s, 5s, 10s, 20s, 30s) and bands (A) delta,(B) theta, (C) alpha, (D) beta, and (E) gamma. Statistical differences were estimated through a bootstrap procedure and significant results are denoted with an asterisk.

### Opinion changes on Scenario 2 time in jail

A considerable amount of subjects (24 subjects) that changed opinions on scenario 2 changed only the time in jail. Since our group division on opinion change binarizes this response, a concern that might arise is if just a small change on the time in jail would classify a subject as having a change in opinion when there actually very little change (< 1 year sentencing change, for example). To address this concern we plot the distribution of differences of time in jail between questionnaire 1 and 2, this is shown in Figure S4. All differences observed accounts for at least 10 years which is a substantial time and was subsequently considered an opinion change.

**Figure S4:**
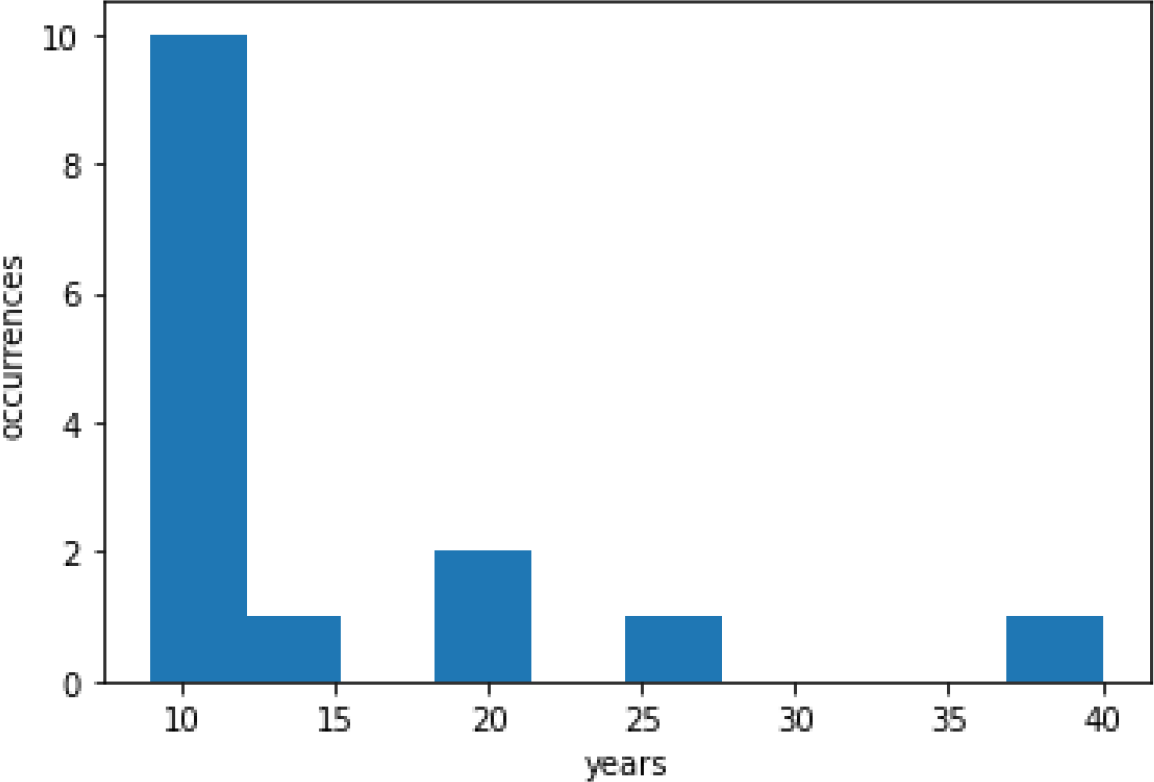
Histogram of time in jail changes in scenario 2. Occurrences of changes on time in jail question after interaction with the social media platform, the minimum change on time observed was 10 years. For this question, 9 subjects that changed their answers opted for either ‘death penalty’ or ‘life sentence’ as an answer in at least one of the two compared questionnaires.

Besides the changes in years in prison for a convicted murder, we also observed 9 subjects that showed opinion changes involving non-numerical values (death penalty and life sentence), the labels for those options were standardized to avoid identification of false differences due to typos and differences on capital letters.

### Example articles presented to subjects

As the posts presented to participants were inspired by real articles, below is a non-exhaustive list of some of content that was linked from the posts, from the travel scenario:

https://electrek.co/2019/04/09/paris-800-electric-buses/

https://www.audleytravel.com/indonesia/country-guides/sulawesi

https://www.bloomberg.com/news/articles/2019-04-06/yellow-vest-protesters-shift-focus-to-paris-business-district

https://www.npr.org/2018/09/28/652489085/strong-quake-hits-along-indonesias-western-sulawesi-island

https://www.express.co.uk/news/world/1110802/france-yellow-vest-protest-paris-riot-emmanuel-macron

https://www.nomadicmatt.com/travel-blogs/how-to-spend-5-days-in-paris/

https://www.youtube.com/watch?v=ErFP51JFUX8

https://www.nytimes.com/2019/03/16/world/europe/france-yellow-vests-protest.html

https://www.nytimes.com/2018/10/02/reader-center/donate-indonesia-tsunami-earthquake-victims.html

### Comparison of link weights using wPLI and PLV

To disentangle the effects of PLV and wPLI, we first inspect the proportion of overlap at a variety of thresholds across wPLI and PLV. Figure S5 below visually depicts the proportion of edges that are overlapping after a binarization of the connectivity matrix for PLV and wPLI. Interestingly, within this dataset, at the extremes of connectivity, where t is the threshold of PLV and wPLI (t < 0.1 || t > 0.8), PLV and wPLI show very similar connectivity patterns; however, at the lower range (t > 0.15 && t < 0.6) there is substantial uniqueness of these connectivity patterns. We conclude that only the mid-range connectivity values have variable amounts of contribution from the common source problem, or volume conduction. In the figure below, we plot the mean, standard deviation, and coefficient of variation across time. For the mean, as the frequency increases, the overall mean decreases relative to the lower frequencies, suggesting that higher frequencies are less likely to be ambiguous in volume conduction effects or the “common sources problem”; however, the mean wPLI of the lower frequencies (e.g., delta, theta, alpha) is within the range of ambiguous sources.

**Figure S5:**
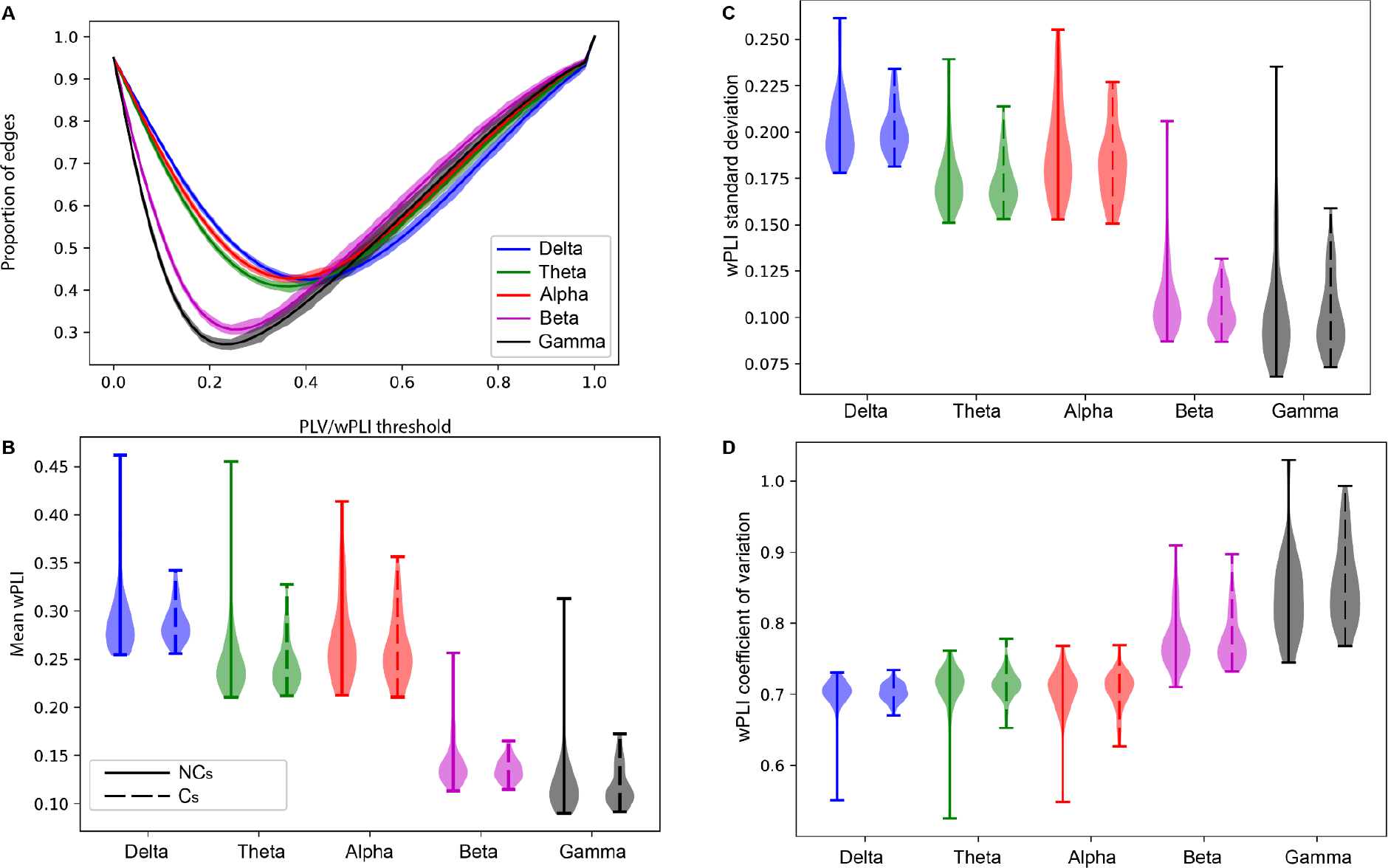
wPLI and PLV differences on the link weights. A Shows the comparison of wPLI and PLV edges present as we increase the threshold for considering an edge as present. B-D show, respectively the mean, standard deviation and coefficient of variation of the wPLI weights for both groups (opinion change and no change). Albeit the fact that the region of the minima is contained in the mean wPLI distribution range, the wPLI coefficient of variation shows that the distribution has a high variance.

**Figure S6:**
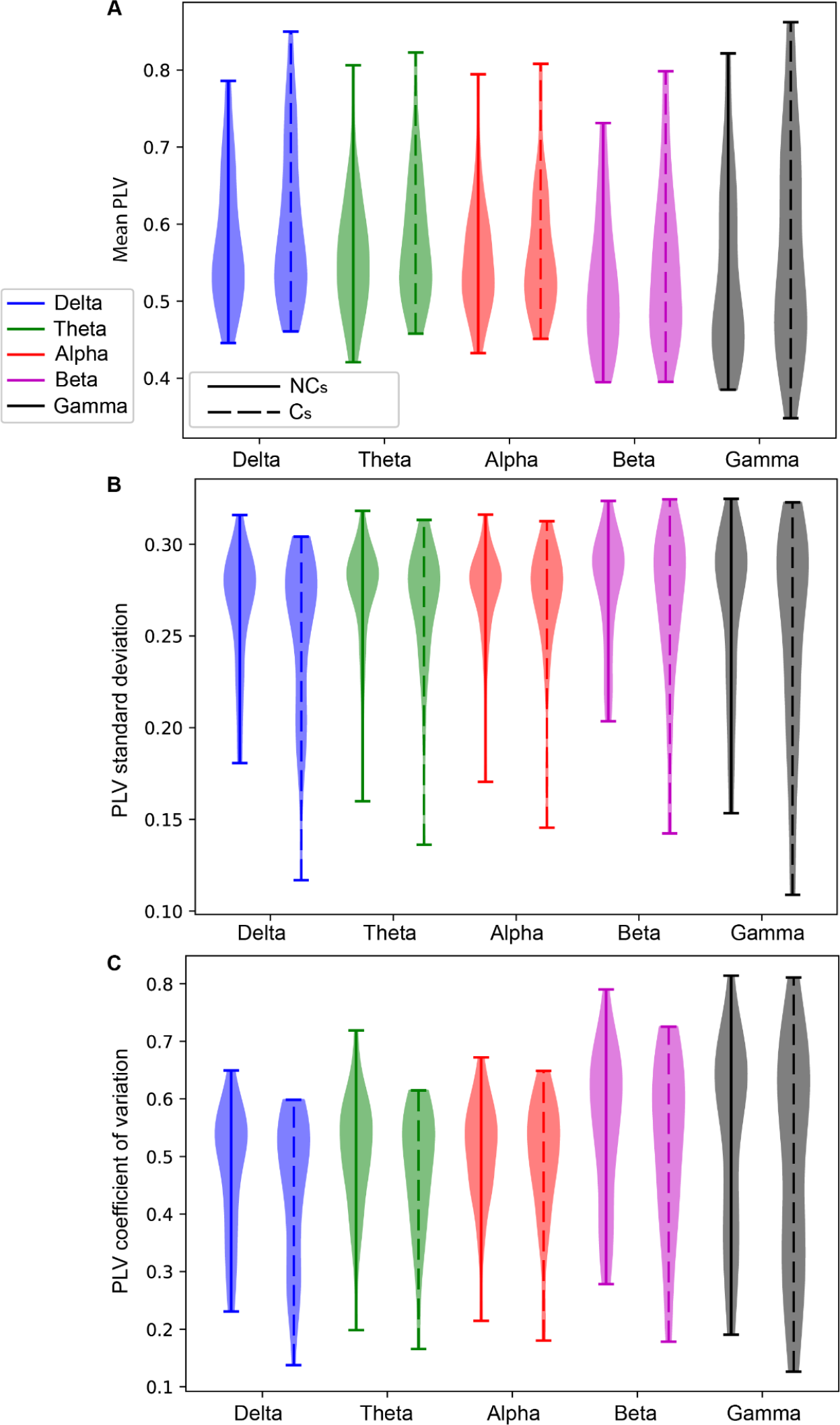
A-C show, respectively the mean, standard deviation and coefficient of variation of the PLV weights for both groups.

The statistics of the weight distributions calculated based on PLV are presented in Figure S6, the distributions cover a larger range of values when compared with wPLI with mean PLV distributed in a range of higher values than wPLI especially for the higher frequency bands. From these two figures, considering the high overlap between metrics at the extreme values and the high variability in PLV that, it is unlikely that volume conduction significantly contributes to our results.

**Figure S7:**
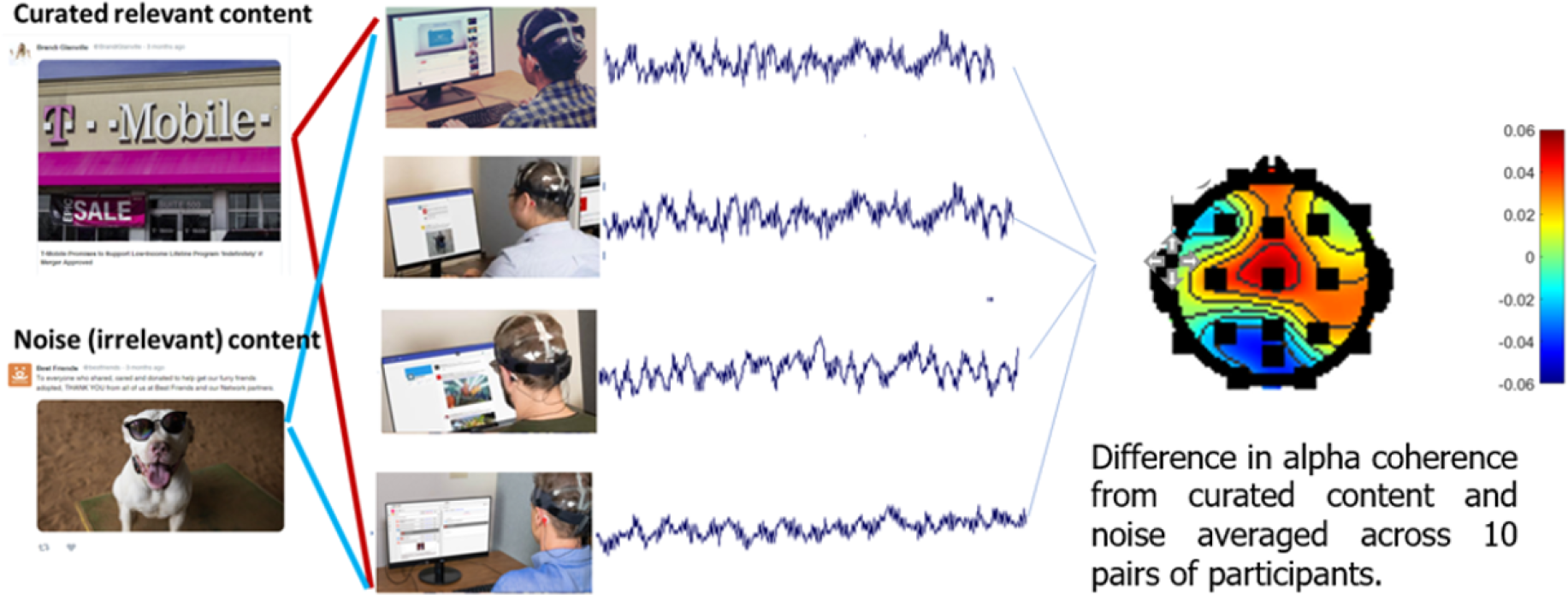
The platform and general setup. Subjects were presented with curated online content to elicit decision making processes (left) whilst instrumented with EEG (middle) and allowed to freely scroll through the content. High level alpha differences between rest and platform engagement are shown (right).

